# Pulmonary dust foci as rat pneumoconiosis lesion induced by titanium dioxide nanoparticles in 13-week inhalation study

**DOI:** 10.1101/2022.03.31.486525

**Authors:** Shotaro Yamano, Yuko Goto, Tomoki Takeda, Shigeyuki Hirai, Yusuke Furukawa, Yoshinori Kikuchi, Tatsuya Kasai, Kyohei Misumi, Masaaki Suzuki, Kenji Takanobu, Hideki Senoh, Misae Saito, Hitomi Kondo, Yumi Umeda

## Abstract

**Background:** Most toxicological studies on titanium dioxide (TiO_2_) particles to date have concentrated on carcinogenicity and acute toxicity, with few studies focusing of pneumoconiosis, which is a variety of airspace and interstitial lung diseases caused by particle-laden macrophages. The present study examined rat pulmonary lesions associated with pneumoconiosis after inhalation exposure to TiO_2_ nanoparticles (NPs).

**Methods:** Male and female F344 rats were exposed to 6.3, 12.5, 25, or 50 mg/m^3^ anatase type TiO_2_ NPs for 6 hours/day, 5 days/week for 13 weeks using a whole-body inhalation exposure system. After the last exposure the rats were euthanized and blood, bronchoalveolar lavage fluid, and all tissues including lungs and mediastinal lymph nodes were collected and subjected to biological and histopathological analyses.

**Results:** Numerous milky white spots were present in the lungs after exposure 25 and 50 mg/m^3^ TiO_2_ NPs. Histopathological analysis revealed that the spots were alveolar lesions, characterized predominantly by the agglomeration of particle-laden macrophages and the presence of reactive alveolar epithelial type 2 cell (AEC2) hyperplasia. We defined this characteristic lesion as pulmonary dust foci (PDF). The PDF is an inflammatory niche, with decreased vascular endothelial cells in the interstitium, and proliferating AEC2 transformed into alveolar epithelial progenitor cells. The AEC2 in the PDF had acquired DNA damage. Based on PDF induction, the lowest observed adverse effect concentration for pulmonary disorders in male and female rats in this study was 12.5 mg/m^3^ and 6.3 mg/m^3^, respectively. The no observed adverse effect concentration for male rats was 6.3 mg/m^3^. There was a sex difference in lung lesion development, with females showing more pronounced lesion parameters than males.

**Conclusions:** Inhalation exposure to TiO_2_ NPs caused PDF, an air-space lesion which is an alveolar inflammatory niche containing particle-laden macrophages and proliferating AEC2. This PDF histopathologically resembles some pneumoconiosis lesions (pulmonary siderosis and hard metal pneumoconiosis) in workers and lung disease in smokers, suggesting that it is an early pneumoconiosis lesion caused by exposure to TiO_2_ NPs in rats and a common alveolar reaction in mammals.

## Background

Titanium dioxide nanoparticles (TiO_2_ NPs) have a variety of applications, from use in sunscreens, toners, and cosmetics to photodynamic therapy and treatment of waste water [1–4]. There are a variety of methods used to synthesize TiO_2_ NPs, resulting in the different particle properties that give TiO_2_ NPs their wide range of applications [1,5,6]. However, extensive use of TiO_2_ NPs without appropriate protections may lead to unexpected effects on human health, such as inhalation toxicity [7–10]. Indeed, there are a number of clinical case reports that workers exposed to TiO_2_ suffered from various lung diseases [11–22]. Based on histopathological similarities, it is suggested that some of these cases include a type of pneumoconiosis, a typical occupational lung disease caused by inhalation of metal dust and fumes [23–26]. Pneumoconiosis is understood histopathologically as a variety of airspace and interstitial lung diseases caused by particle-laden macrophages, which are chronic, progressive and still have no fundamental treatment [23–26]. Furthermore, the progression of pneumoconiosis is well known to complicate lung cancer [27, 28]. Therefore, there is an urgent need to understand the toxicity mechanisms and pathogenesis of TiO_2_-associated pneumoconiosis to safe-guard the health of workers handling TiO_2_ NPs. However, most toxicological studies on TiO_2_ particles to date have focused on carcinogenicity and acute toxicity, and few studies have investigated the development of pneumoconiosis.

Although not a study focused on pneumoconiosis, one subchronic inhalation toxicity study of TiO_2_ NPs (P25 containing both anatase and rutile forms of TiO_2_ obtained from DeGussa-Huls AG, mean primary particle size 21 nm) has been conducted using rats, mice, and hamsters [29]. In this report, it was found that inhalation of 10 mg/m^3^ TiO_2_ NP for 13 weeks caused inflammatory responses in both rats and mice. In addition, similarly to humans exposed to TiO_2_ [22], rats developed progressive fibroproliferative lesions with interstitial particle accumulation, and alveolar septal fibrosis. These findings suggest that 13 weeks of inhalation exposure to rats is a sufficient duration to observe progressive lung lesions caused by TiO_2_ NPs and is an appropriate experimental protocol to assess the early stages of pneumoconiosis.

The P25 TiO_2_ NPs used in the study by Bermudez et al. contained a mixture of two types of TiO_2_ crystal structures, anatase and rutile; therefore, the toxicities due to anatase TiO_2_ and rutile TiO_2_ were not distinguished from one another. In the present study, male and female rats were exposed to anatase TiO_2_ NPs for 13 weeks by systemic inhalation to investigate dose-response pathological changes associated with pneumoconiosis and to define the histopathological and cell biological basis of the development of pulmonary lesions associated with pneumoconiosis.

Adverse outcome pathways (AOPs) are conceptual frameworks that link initiating events (IEs) and adverse outcomes (AOs) via key events (KEs) using known information and has been used to understand how certain adverse events occur [30]. Although there is no AOP for pneumoconiosis, a putative AOP for carcinogenicity resulting from inhalation of TiO_2_ has been summarized in a previous report [31]. Briefly, particle-laden macrophages remain in the alveolar airspace, causing impaired clearance of the inhaled particles and persistent inflammation (KE3). Persistent inflammation leads to persistent tissue damage (persistent epithelial injury: KE4) and repair (epithelial cell proliferation: KE6). Alveolar epithelial cell (AEC) proliferation in the presence of inflammatory agents can result in genetic damage in the replicating cells (KE5), resulting in DNA mutations being passed to the daughter cells. Continued tissue damage and repair leads to continued proliferation of AEC containing DNA mutations, allowing these cells to acquire additional mutations and to expand (KE6), resulting in bronchiolo-alveolar hyperplasia containing epithelial cells with genetic damage, a pre-neoplastic epithelial lesion (KE7). Lung fibrosis, which is also a feature of pneumoconiosis, has been discussed as a KE in the inhalation toxicity of NPs containing TiO_2_ [32]. Therefore, in addition to assessing pulmonary lesions associated with pneumoconiosis, we examined the lung lesions caused by inhalation exposure to TiO_2_ NPs for the presence of the KEs associated with tumorigenesis mentioned above and for the presence of fibrosis.

## Results

### Stability of aerosol generation and mass concentration and particle size distribution of TiO_2_ NPs in the inhalation chamber

The mass concentrations of TiO_2_ NPs aerosol in the inhalation chambers are shown in Figure S1. Each TiO_2_ NPs concentration was nearly equal to the target concentration the 13-week exposure period (Fig. S1B). The size distribution and morphology of the particles was measured at the first, sixth, and last week of exposure. The mass median aerodynamic diameters (MMAD) and geometric standard deviations (σg) of the TiO_2_ aerosols were within 0.9-1.0 μm and 2.0-2.1, respectively, and were similar for all TiO_2_ NPs-exposed groups (Fig. S1D). Morphological observations by scanning electron microscope (SEM) confirmed that the TiO_2_ NPs generated in the chamber did not appear to be highly aggregated (Fig. S1E). These data indicate that the size distribution and morphology of the TiO_2_ NPs aerosols were consistent during the 13-week exposure period.

### Final body weights and organ weights

In all TiO_2_ NP-exposed rats, neither exposure-related mortality nor respiratory clinical signs were observed throughout the study. There were no significant changes in final body weight (Fig. 1A, B). TiO_2_ NP concentration-dependent increases in lung weight were observed in both males and females (Fig. 1C-F). No statistically significant changes in the weight of organs other than the lungs were observed in any of the exposure groups (Tables S1 and S2).

**Figure 1.**
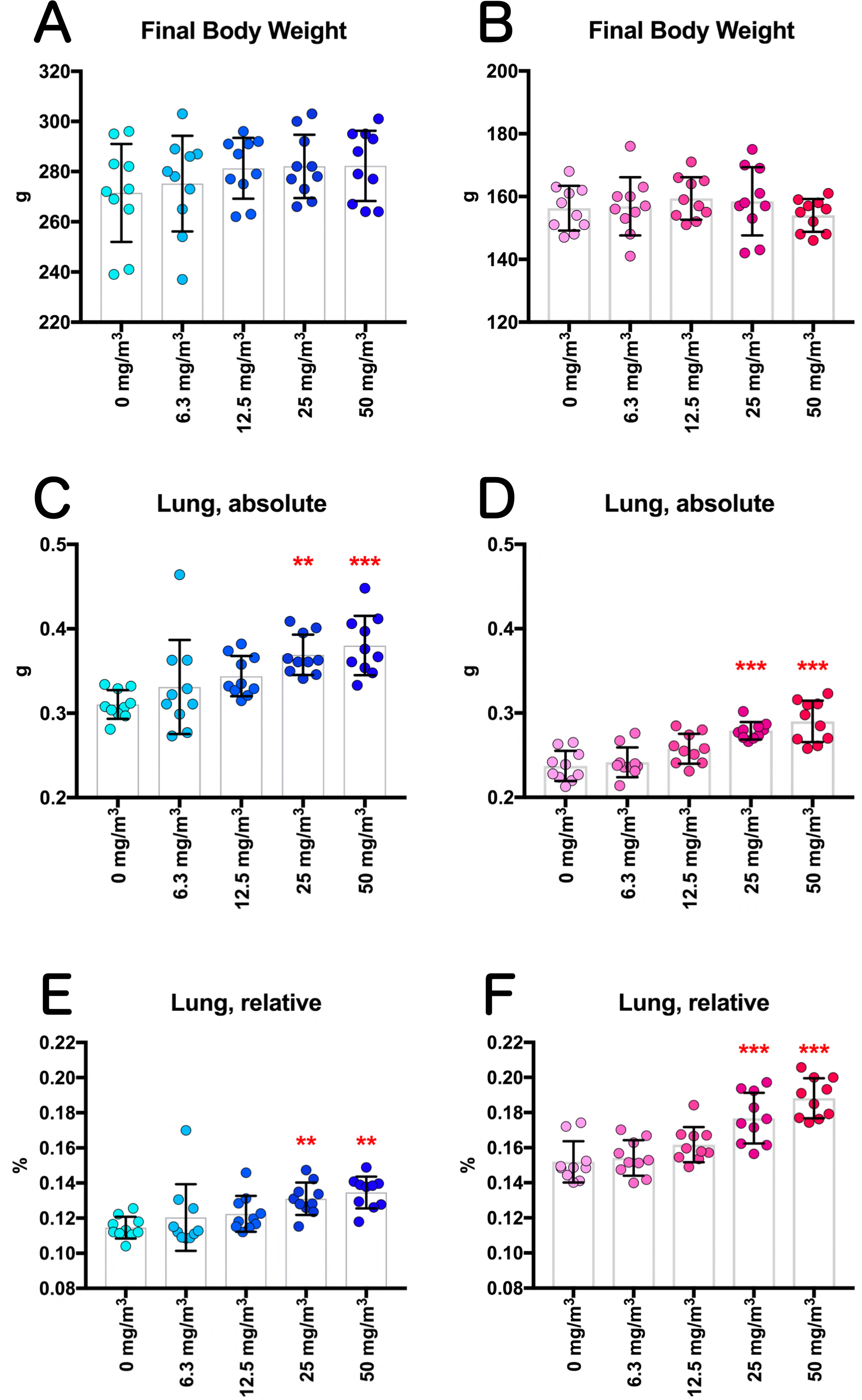
Final body weights and lung weights of F344 rats exposed to titanium dioxide nanoparticles (TiO_2_ NPs) by inhalation (6.3, 12.5, 25, or 50 mg/m^3^, 6 hours/day, 5days/week, 13 weeks). Final body weights of male (A) and female (B) rats. Absolute left lung weights in male (C) and female (D) rats. The relative lung weights in male (E) and female (F) rats were calculated as a percentage of body weight. Dunnett’s multiple comparison test: \*\**p*<0.01, and \*\*\**p*<0.001.

### Blood hematology and biochemistry

Blood hematology and biochemistry data is shown in Tables S3 and S4. A significant increase in the percentage of eosinophils in the white blood cells (WBCs) was observed in males exposed to 12.5 mg/m^3^ and higher concentrations of TiO_2_ (Table S3), and plasma lactate dehydrogenase (LDH) and aspartate aminotransferase (AST) activity and urea nitrogen levels were significantly increased in females in the 50 mg/m^3^ exposure group (Table S4). However, these were judged to be low toxicological significance, because the changes were small and did not occur in both males and females.

### Measurement of cytological and biochemical markers in the bronchoalveolar lavage fluid (BALF)

In the clean air groups of both sexes, normal macrophages with fine vacuoles were predominantly observed in the BALF (Fig. 2). However, in the BALF of both male and female 50 mg/m^3^ groups, a number of neutrophils and enlarged macrophages phagocytosing TiO_2_ NP were observed (Fig. 2). Cell population analysis found that the total cell number in the BALF of the male and female 50 mg/m^3^ exposure groups was significantly increased (Fig. 3A and B). Neutrophil numbers increased in a TiO_2_ NP concentration-dependent manner and were significantly increased in males exposed to 25 and 50 mg/m^3^ and in females exposed to 50 mg/m^3^ TiO_2_ (Fig. 3C and D). Lymphocyte numbers increased in a TiO_2_ NP concentration-dependent manner and were significantly increased in male and female 50 mg/m^3^exposure groups (Fig. 3E and F). In contrast, there was no increase in alveolar macrophage (AM) numbers in any of the exposure groups (Fig. 3G and H).

**Figure 2.**
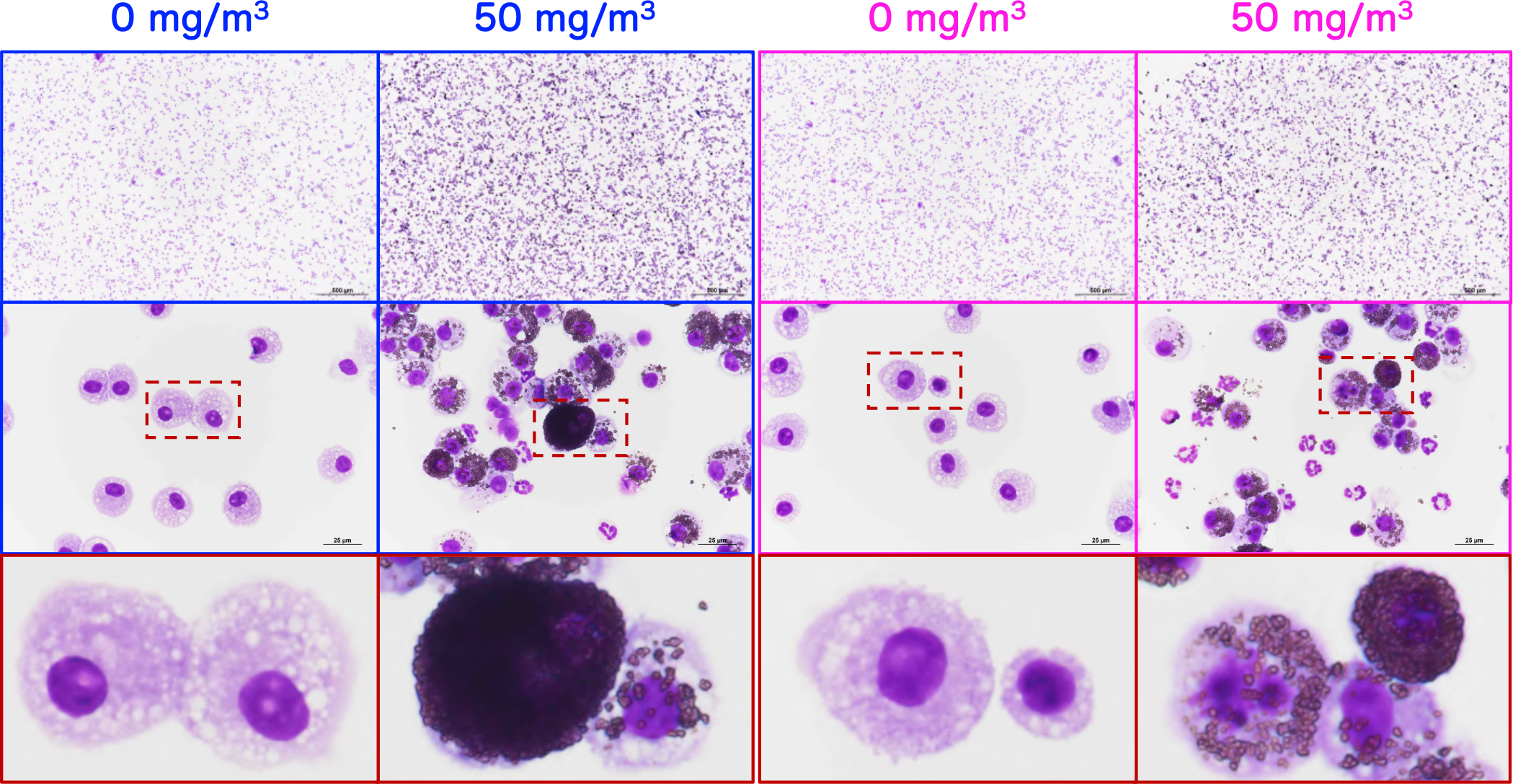
Representative images of the bronchoalveolar lavage fluid (BALF) cytospin cytology. The BALF samples obtained from males (blue) and females (pink) exposed to 0 mg/m^3^ or 50 mg/m^3^ TiO_2_ NPs were stained with May-Grunwald-Giemsa. The area enclosed by the red dotted square in the middle panel is enlarged and shown in the bottom panel.

**Figure 3.**
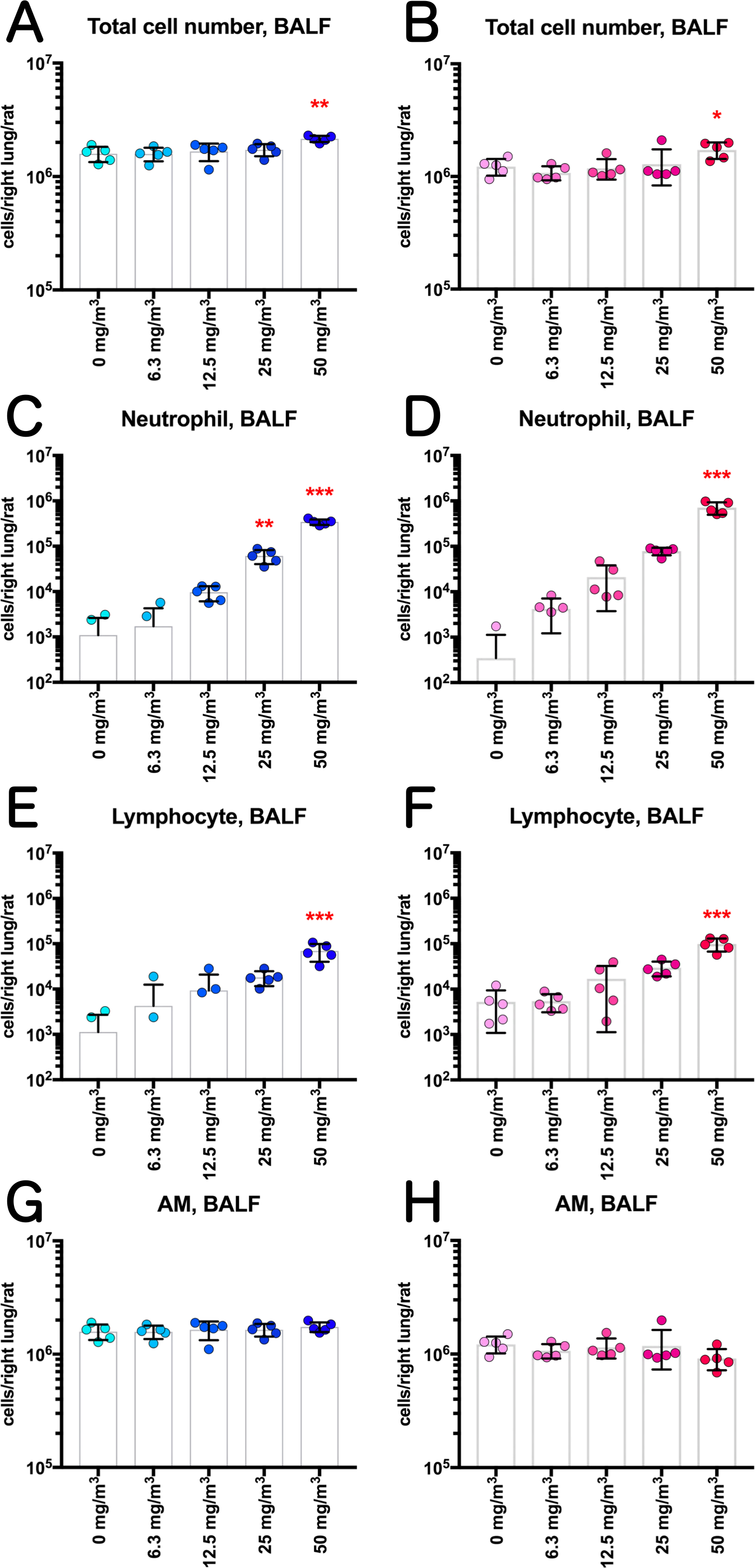
Effect of inhalation exposure to TiO_2_ NP on cell number in the BALF. The number of total cells (A, B), neutrophils (C, D), lymphocytes (E, F), and alveolar macrophages (AM) (G, H) were counted using an automated hematology analyzer, and are shown by sex (males: A, C, E and G; females: B, D, F and H). Statistical significance was analyzed using Dunnett’s multiple comparison test: \**p*<0.05, \*\**p*<0.01, and \*\*\**p*<0.001.

TiO_2_ NP exposure increased LDH activity and total protein and albumin levels, but not alkaline phosphatase (ALP) or γ-glutamyl transpeptidase (γ-GTP) activities, in the BALF (Figs. 4 and S2). In males, these increases showed clear concentration dependence. In females, significant increases were only observed in the 50 mg/m^3^ group, however, in the 50 mg/m^3^ groups females showed more pronounced increases than males.

**Figure 4.**
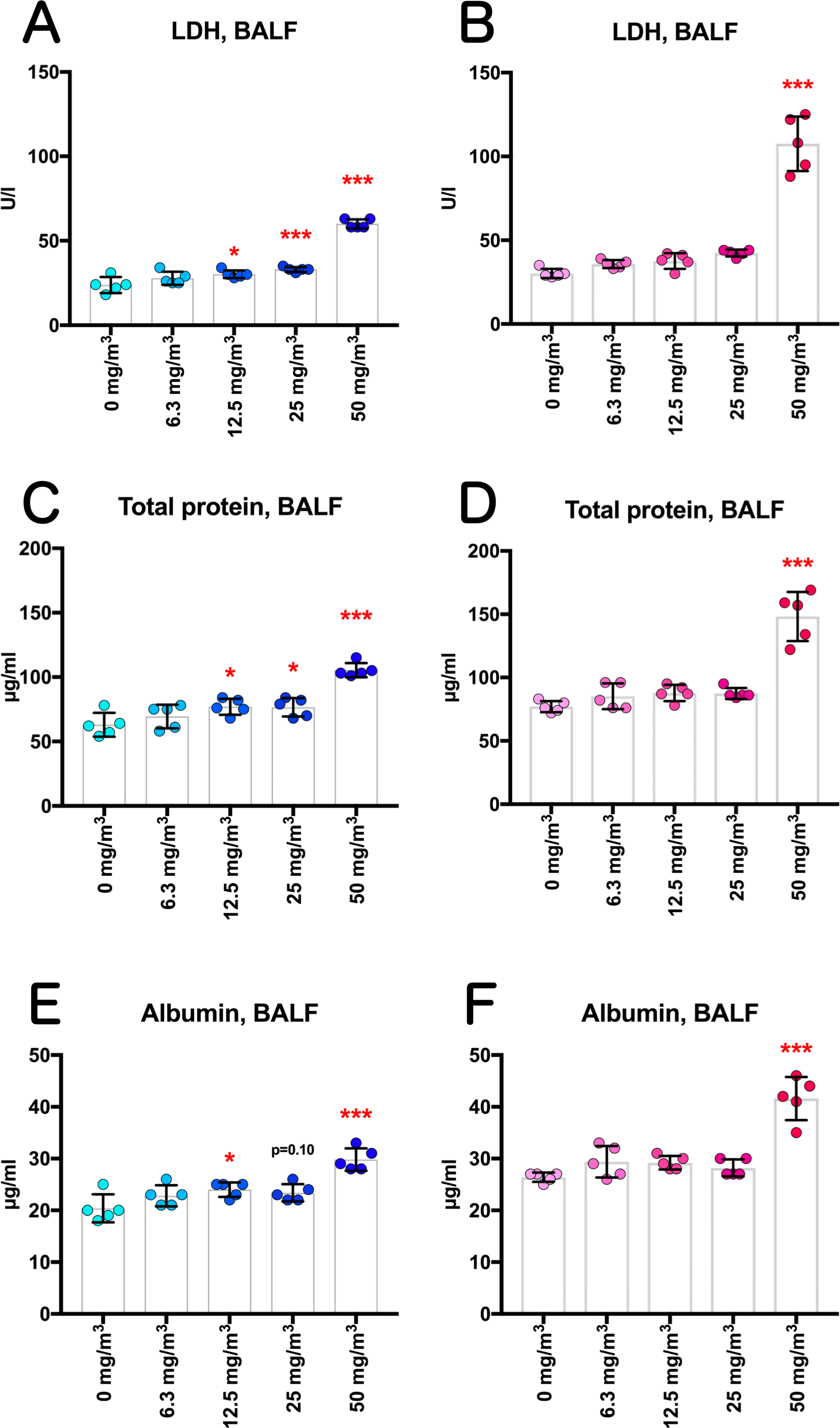
Dose-dependent induction of biochemical markers in the BALF obtained from the lungs of rats after inhalation of TiO_2_ NP for 13 weeks. Lactate dehydrogenase (LDH) activity (A, B), total protein concentration (C, D), and albumin concentration (E, F) in the BALF were measured using an automatic analyzer, and are shown by sex (males: A, C and E; females: B, D and F). Statistical significance was analyzed using Dunnett’s multiple comparison test: \**p*<0.05, and \*\*\**p*<0.001.

BALF cytospin specimens revealed that AMs were present in various TiO_2_ NPs-phagocytic states (Fig. 5A). AMs that phagocytosed one or more TiO_2_ NPs were defined as TiO_2_ NP-laden AMs. TiO_2_ NP-laden AMs that phagocytosed TiO_2_ NPs until the nucleus was no longer visible were defined as Over-stuffed, and AMs that disintegrated into particles and cellular debris were defined as Burst (Fig. 5A). On average, more than 99% of AMs in all exposure groups of both sexes were TiO_2_ NP-laden (Fig. 5B and C). The percentage of Over-stuffed AMs increased in both sexes in an exposure concentration-dependent manner, with a statistically significant increase in the groups exposed to 25 and 50 mg/m^3^ (Fig. 5D and E). There was no concentration-dependent increase in the percentage of Burst AMs in either sex (Fig. 5F and G).

**Figure 5.**
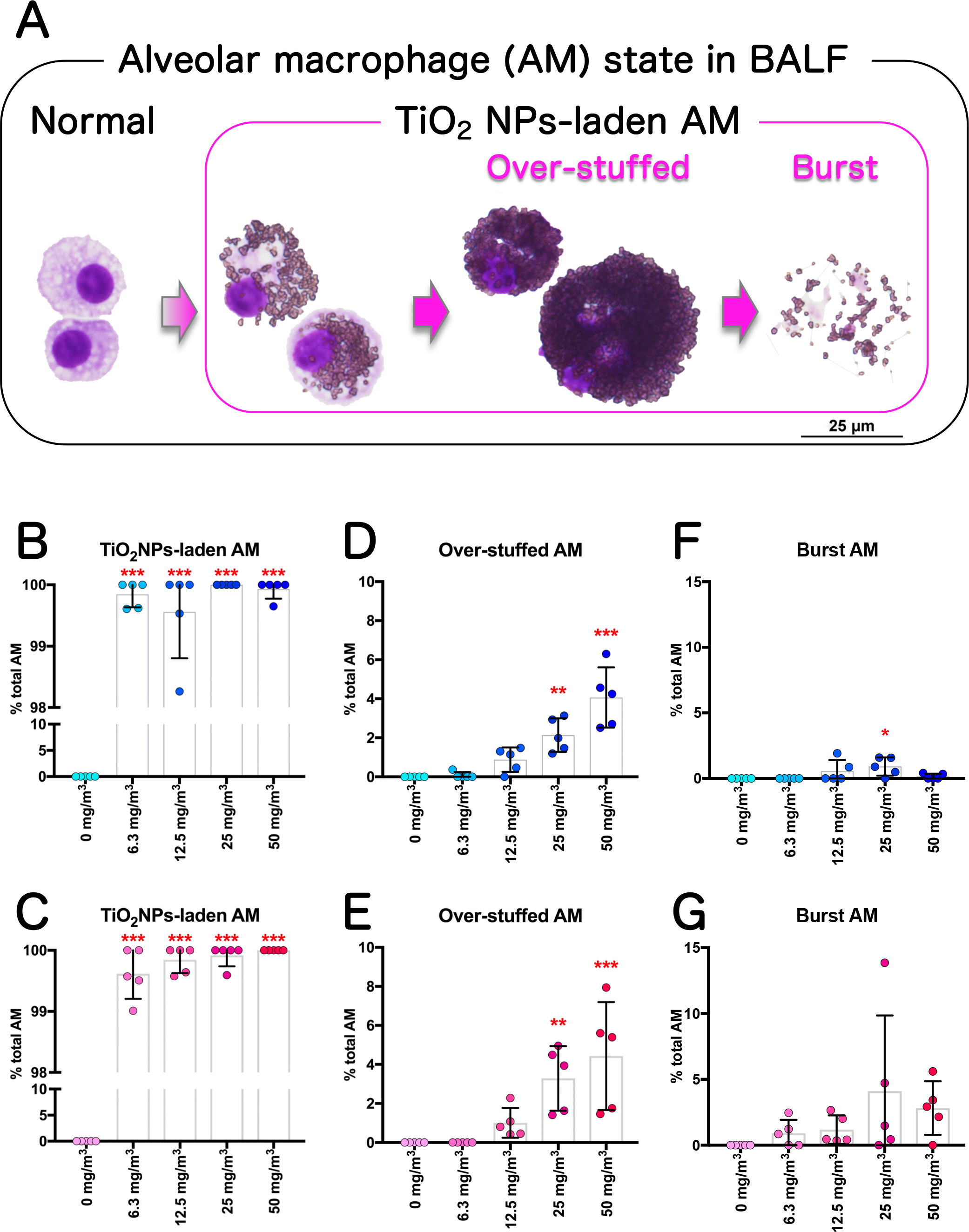
Additional analyses of alveolar macrophages (AMs) in BALF cytospin cytology. Various states of AMs phagocytosing TiO_2_ NPs were found by careful observation of BALF cytospin specimens (A). AM that phagocytosed one or more TiO_2_ NPs were defined as TiO_2_ NP-laden AMs. AM that phagocytosed TiO_2_ NPs until the nucleus was no longer visible were defined as Over-stuffed AM. AM that disintegrated into particles and cellular debris were defined as Burst AM. The percentage of TiO_2_ NP-laden AM (B, C), Over-stuffed AM (D, E), and Burst AM (F, G) were counted, and are shown by sex (males: B, D and F; females: C, E and G). Statistical significance was analyzed using Dunnett’s multiple comparison test: \*\**p*<0.01, and \*\*\**p*<0.001.

### Lung burden and its correlation with lung weight and BALF markers

OECD TG 413 recommends that lung burden should be measured in a range-finding study or when inhaled test particles are poorly soluble and are likely to be retained in the lungs [33]. Therefore, we investigated the correlation of lung burden with toxicological parameters. Lung burden measurements are shown in Fig. 6A and 6B. Inhalation of TiO_2_ NP resulted in deposition of particles in the lungs in an exposure concentration-dependent manner, which tended to be higher in females than males in the 50 mg/m^3^ group (Fig. 6A and B). Relative lung weight, LDH activity in the BALF, and neutrophil count in the BALF were all positively correlated with lung burden in both sexes (Fig. 6C-H). The correlation plots of lung burden and LDH and lung burden and neutrophil count (Fig. 6E-H) appear to divide into two clusters: data points from the 6.3, 12.5, and 25 mg/m^3^ exposed rats and data points from the 50 mg/m^3^ exposed rats.

**Figure 6.**
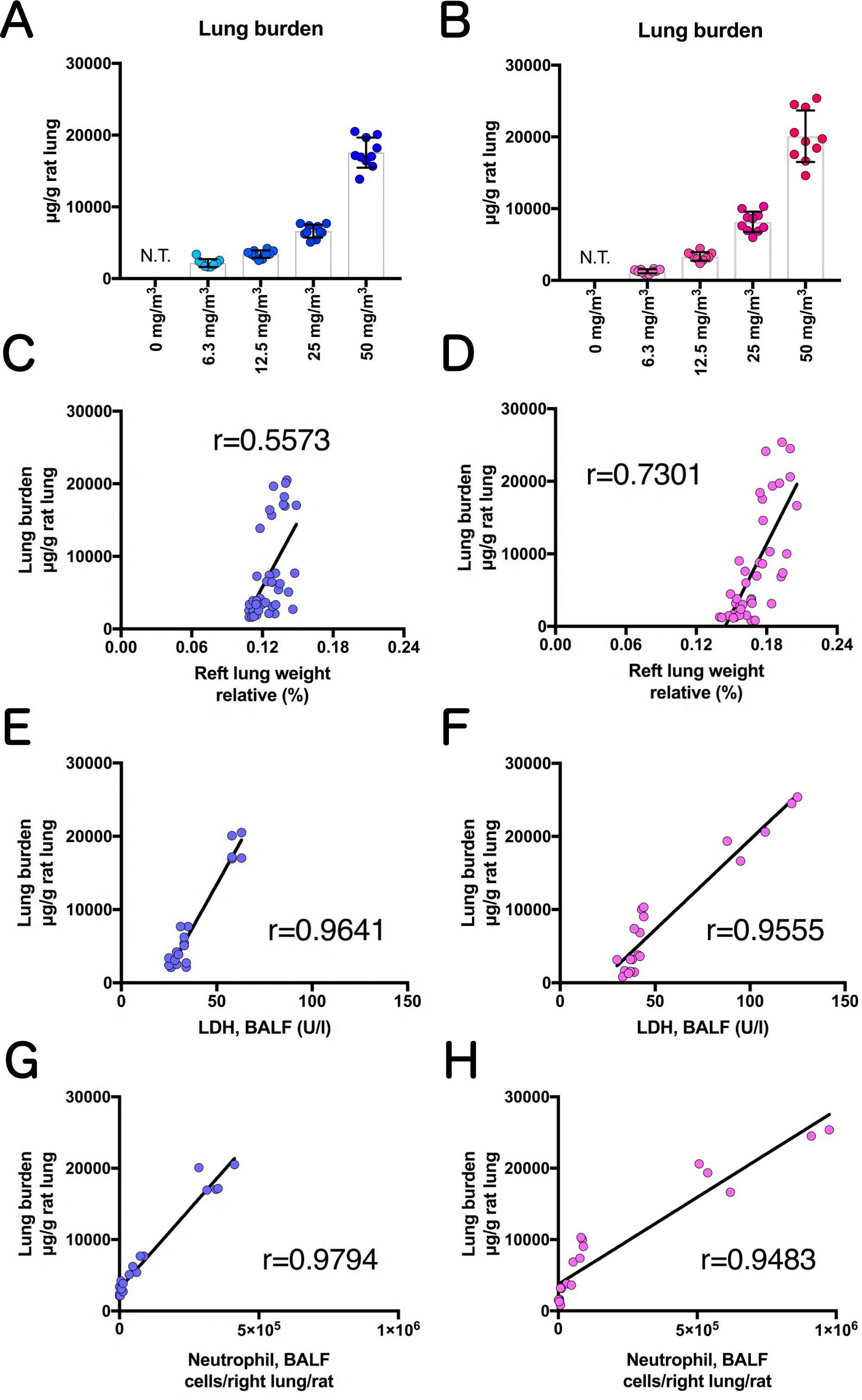
Lung burden and correlation between lung burden and relative lung weights or BALF markers. Lung burden of TiO_2_ NPs in male (A) and female (B) rats was measured by a Zeeman atomic absorption spectrometry. Correlation between lung burden and relative lung weight (C, D), LDH activity in the BALF (E, F), or neutrophil number in the BALF (G, H) was analyzed using the Pearson’s correlation coefficients, and are shown by sex (male: C, E and G; female: D, F and H). r: Pearson’s correlation coefficient. Abbreviation: N.T., not tested.

### Macroscopic findings of lung and mediastinal lymph node

Representative macroscopic images of the lungs and mediastinal lymph nodes are shown in Figs. 7 and S3. In the lungs of the 25 and 50 mg/m^3^ exposed rats, a large number of milky white spots were observed in all lung lobes, mainly on the lung surfaces facing the ribs. The spots were generally approximately 300 nm in diameter, but some were partially fused and were about 1 mm in diameter (Fig. 7). The spots observed on the lung surface were fewer around the hilar region and more concentrated at the lung periphery (Fig. S3B). The mediastinal lymph nodes also showed a similar color change (Fig. 7). However, no significant enlargement of mediastinal lymph nodes due to TiO_2_ NP exposure was observed. These gross changes in the lungs and mediastinal lymph nodes were observed only in the groups exposed to 25 and 50 mg/m^3^.

**Figure 7.**
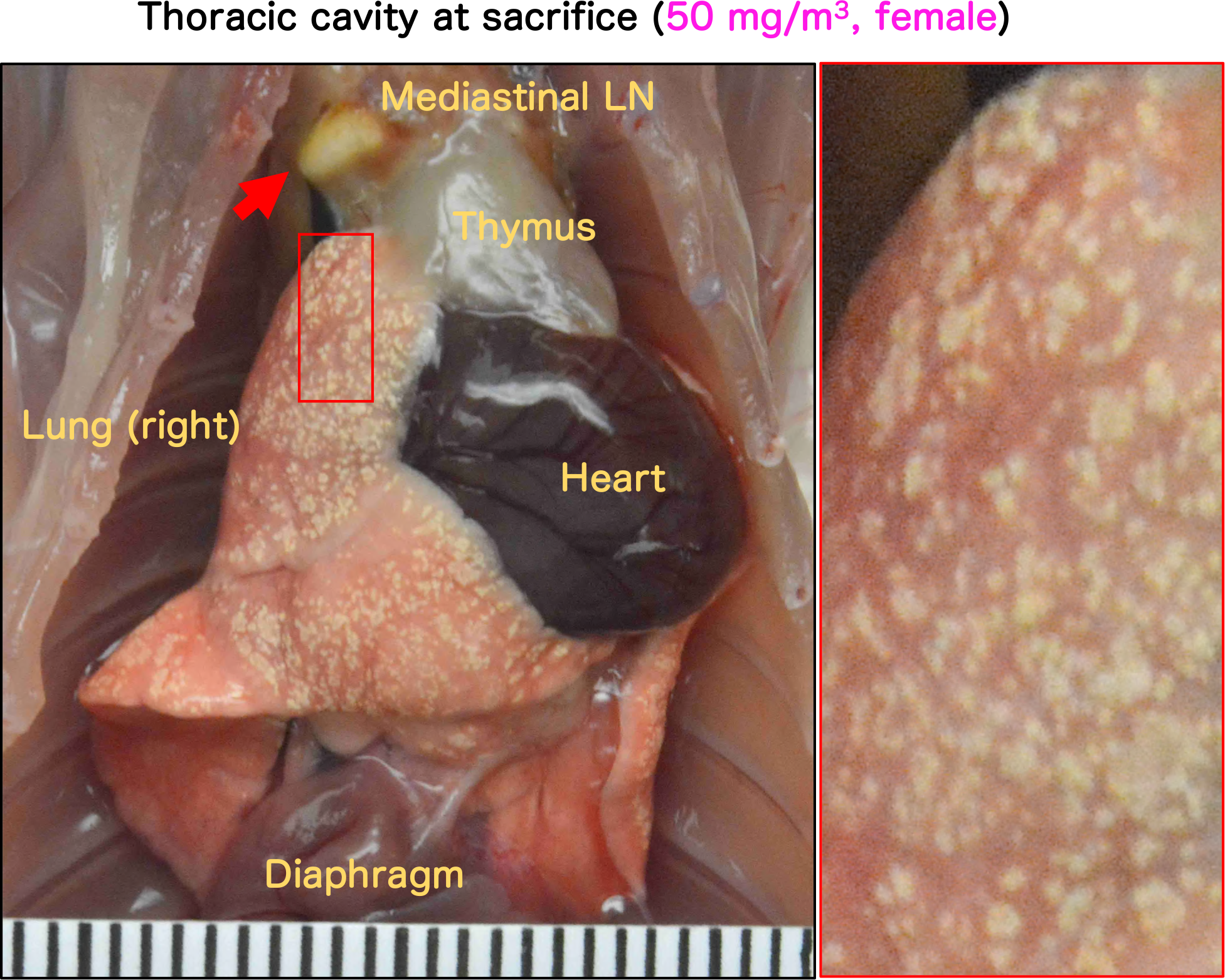
Representative macroscopic photographs of the thoracic cavity of female rats after 13-week of inhalation exposure to TiO_2_ NP (50 mg/m^3^, 6 hours/day, 5 days/week). Abbreviation: LN, lymph node

### Histopathological examination for lung and mediastinal lymph node

Representative microscopic photographs and histopathological findings for the lung and mediastinal lymph nodes are shown in Figs. 8, S4, S5, S6 and Table 1. TiO_2_ NP exposure induced various particle-laden macrophage-associated pulmonary lesions. Deposition of particles in the alveolar air space (Fig. 8B), bronchus-associated lymphoid tissue (BALT) (Fig. S4C), and mediastinal lymph node (Fig. S6), commonly seen with inhalation exposure to particles, was observed in all groups exposed to TiO_2_ NPs. The lesion intensity was concentration-dependent (Table 1). Extrapulmonary ejection of TiO_2_ NPs via the mucociliary escalator was observed in the all exposed groups (Fig. S4B).

**Figure 8.**
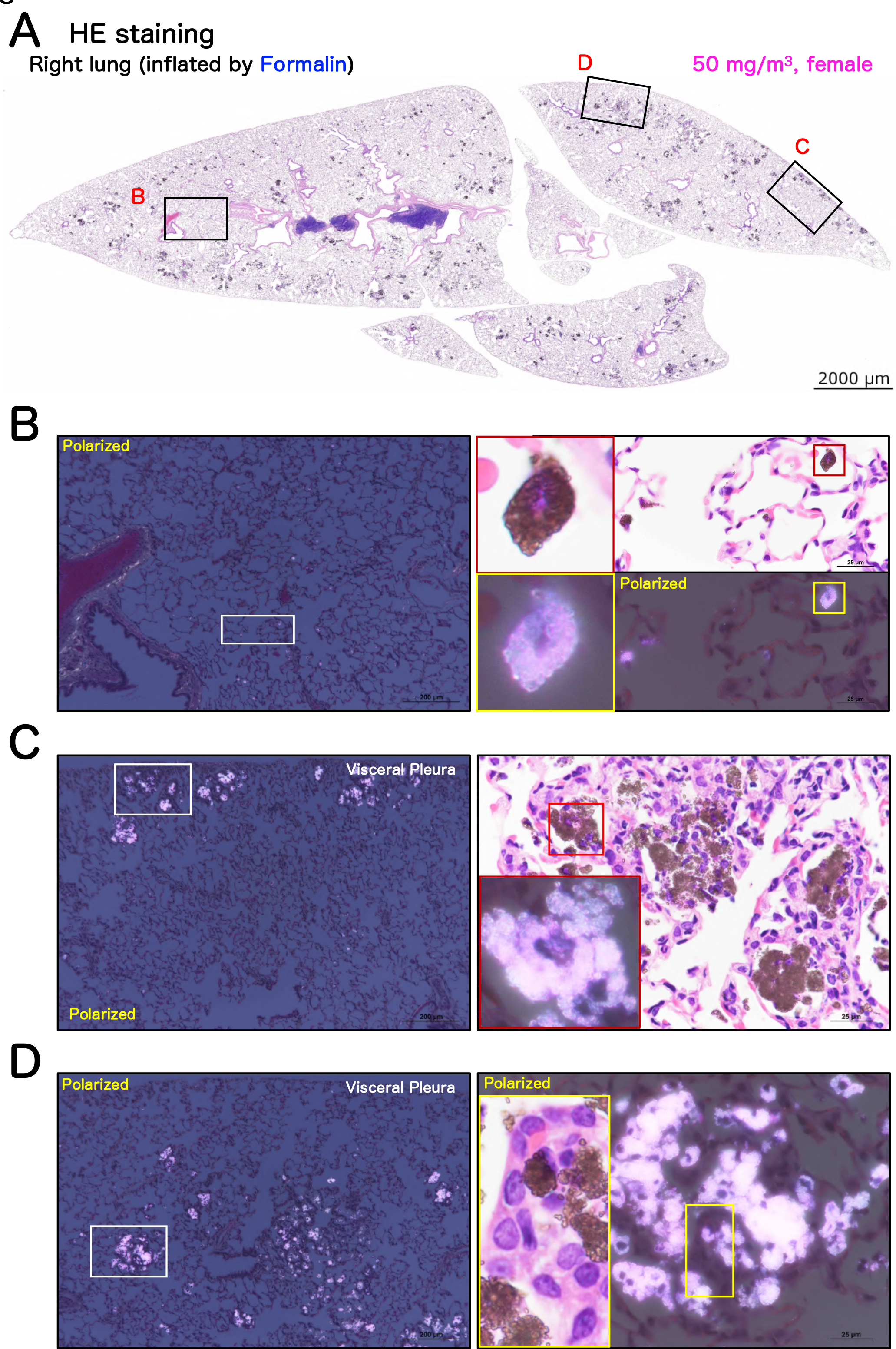
Representative microscopic photographs of female rat lungs after inhalation exposure to TiO_2_ NP (50 mg/m^3^). Representative loupe photograph (A) and magnified images of the area surrounding a lesion (B) and TiO_2_ NP-induced multifocal lesions (C, D) are shown. Formalin was injected into the right lung through the bronchus and the tissue was stained with hematoxylin and eosin (HE). All images were taken with HE or a polarized light microscope. TiO_2_ NPs phagocytosed by macrophages were observed throughout the alveolar region and were black in HE staining or pink in polarized light microscope. Macrophages phagocytosing particles until the nucleus was not visible (magnified right side of panel B). Representative histological images of multifocal lesions in the area just below the pleura (C) and around the Bronchiolo-alverolar duct junction (D) are shown. Histopathologically, in common with each lesion, particle-laden macrophage agglomeration was observed, and in many foci, alveolar epithelial proliferation and inflammatory cell infiltration was also observed (magnified right side of panels C and D).

**Table 1.**
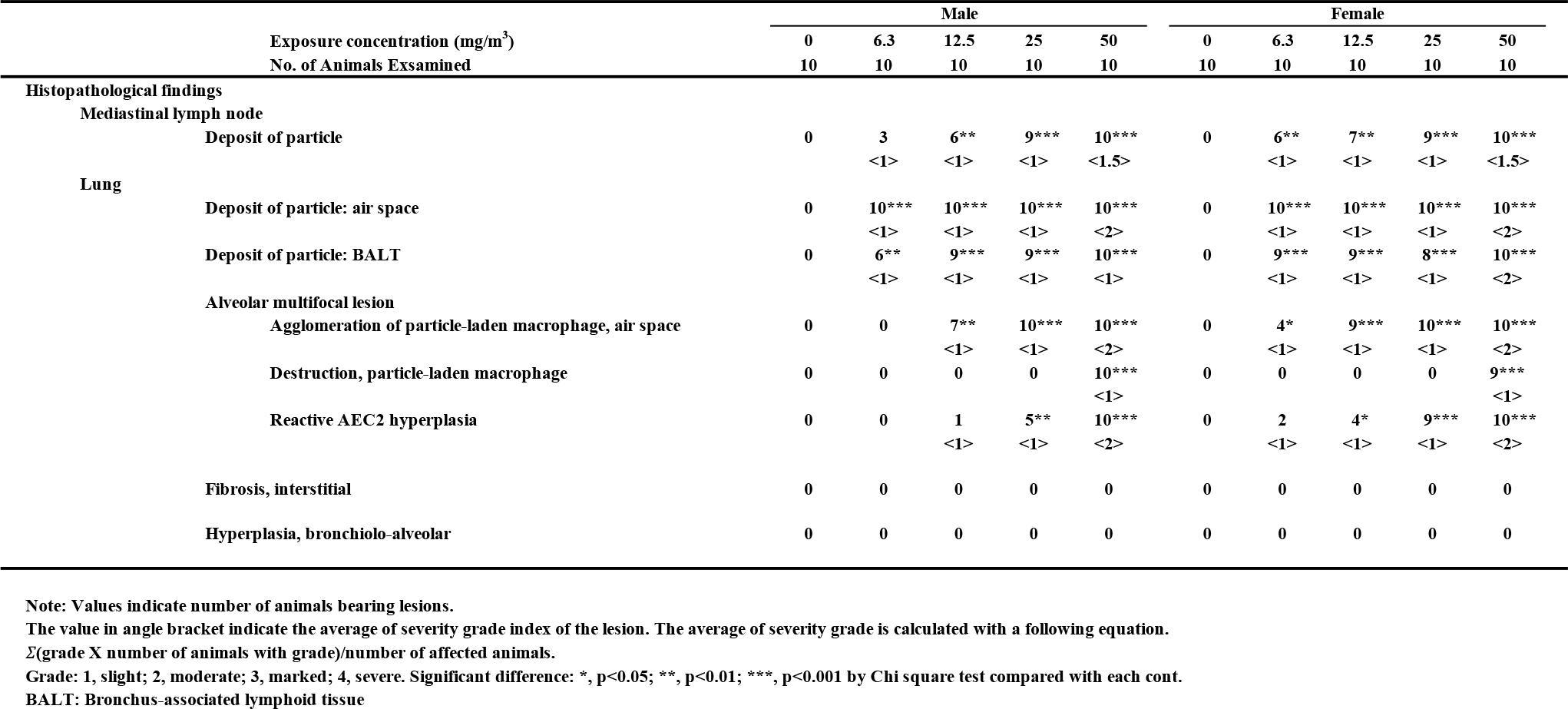
Incidence and integrity of the histopathological findings of the lung and mediastinal lymph nodes after inhalation exposure to TiO_2_ NP.

The milky white spots on the lung surface observed by macroscopic observation were histopathologically identified as multifocal lesions consisting predominantly of agglomerations of particle-laden macrophages in the alveolar air spaces, which were observed as black areas by HE staining (Figs. 8A and S4A) and as pink areas by polarized light (Figs. 8C, 8D, S4B, S4C and S4D). These multifocal lesions of the alveoli were located in the peripheral subpleural area (Fig. 8C), or in the alveolar region around the terminal bronchioles in the hilar region (Fig. 8D). Particle-laden macrophages in the multifocal lesions phagocytosed TiO_2_ NP to the extent that the nuclei were obscured, similar to the “Over-stuffed AM” observed in the BALF (Fig. 8C and D). Macrophages that had burst open, releasing their contents, were also observed in these multifocal lesions (Fig. S4D). Similarly, the neutrophils and lymphocytes observed in the BALF were mainly observed within the multifocal lesions, suggesting that these multifocal lesions are inflammatory niches.

Notably, proliferative changes in the alveolar epithelium were present in the multifocal lesions (magnified view on the right in Fig. 8D). To confirm the cell types constituting the multifocal lesions, multiple staining with cell-specific markers was performed (Figs. 9, S7, S8 and S9). The results showed that all the epithelial cells in the lesions were negative for the club cell marker club cell secretory protein (CCSP), the neuroendocrine cell marker calcitonin gene-related peptide (CGRP), the basal cell marker p63, and the bronchial epithelial lineage marker SRY-Box Transcription Factor 2 (Sox2) (Fig. S7B). In contrast, the alveolar epithelium proliferating in multifocal lesions was positive for the alveolar epithelial type 2 cell (AEC2) marker lysophosphatidylcholine acyltransferase 1 (LPCAT1), indicating that AEC2 hyperplasia is co-localized with agglomeration of particle-laden macrophages in multifocal lesions (Fig. 9A). We defined this type of multifocal lesions containing AEC2 hyperplasia as “Pulmonary dust foci (PDF)”, and distinguished them from “Agglomeration” lesions that do not contain AEC2 hyperplasia.

**Figure 9.**
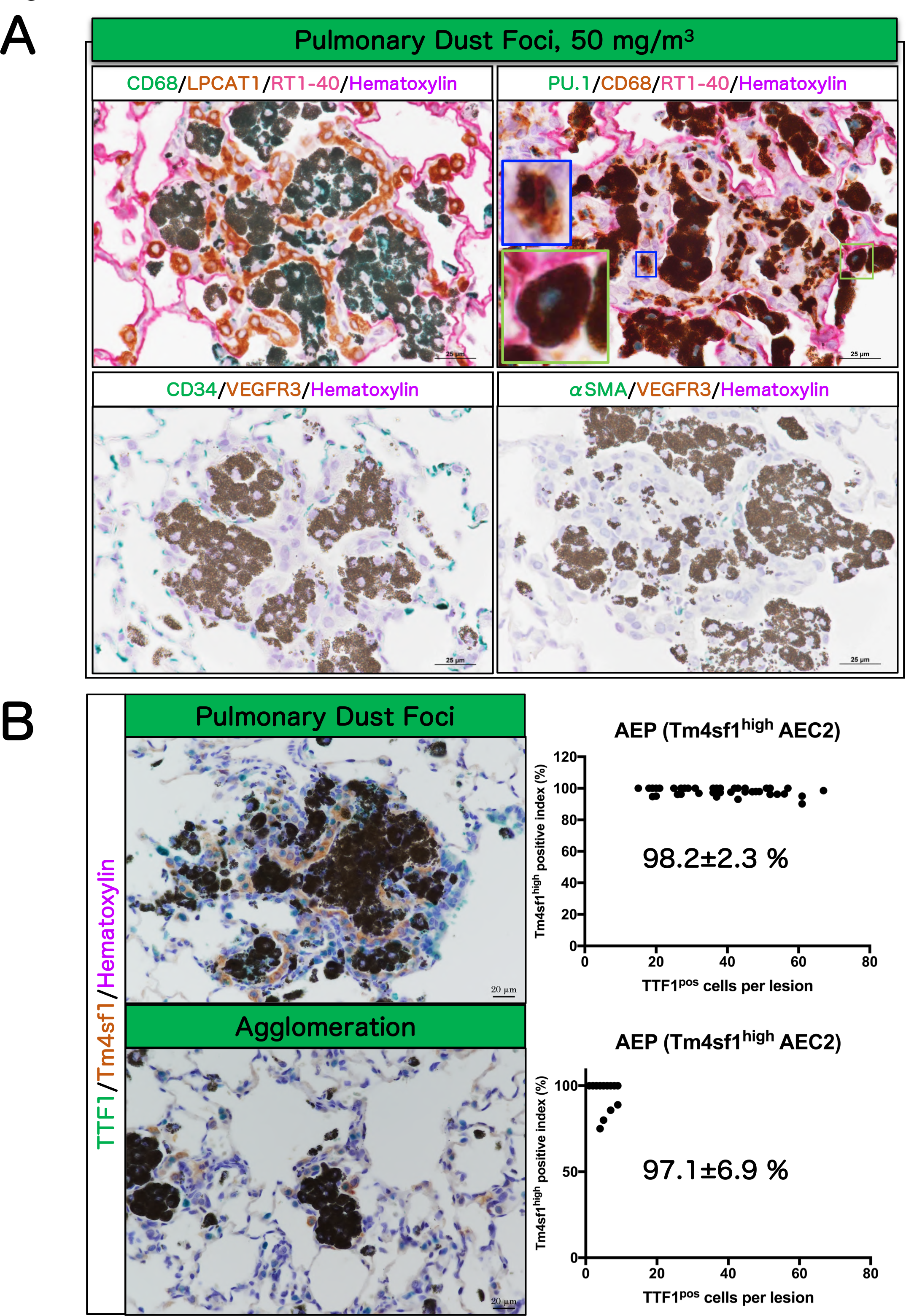
Immunohistochemical characteristics of pulmonary dust foci (PDF) and agglomeration in rat lungs after inhalation exposure to TiO_2_ NP (50 mg/m^3^). We defined multifocal lesions by distinguishing between “Pulmonary dust foci (PDF)”, which are lesions associated with alveolar epithelial type 2 cell (AEC2) hyperplasia, and “Agglomeration”, which are lesions without AEC2 hyperplasia. Triple staining for set 1 (macrophage marker CD68: green in the cytoplasm; AEC2 marker lysophosphatidylcholine acyltransferase (LPCAT) 1: brown in the cytoplasm; AEC1 marker RT1-40: red in the cell membrane; upper left of A). Triple staining for set 2 (myeloid lineage marker PU.1: green in the nucleus; macrophage marker CD68: brown in the cytoplasm; AEC1 marker RT1-40: red in the cell membrane; upper right of A). Double staining for set 3 (vascular endothelial cell marker CD34: green in the cell membrane; lymphatic endothelial cell marker vascular endothelial growth factor receptor (VEGFR) 3: brown in the cell membrane; lower left of A). Double staining for set 4 (myofibroblast marker α-smooth muscle actin (αSMA): green in the cytoplasm; lymphatic endothelial cell marker VEGFR3: brown in the cell membrane; lower right of A). Blue frame in right of A: Particle-laden interstitial macrophage. Green frame in right of A: Over-stuffed AM in the alveolar air space. We have previously reported that Tm4sf1-positive AEC2 (alveolar epithelial progenitor, AEP) appear during the regeneration process after lung injury [30, 31]. Double staining for both PDF and agglomeration (AEC2 marker TTF1: green in the nucleus; AEP marker Tm4sf1: brown in the cytoplasm) (B). The percentage of TTF1/Tm4sf1 double positive AEPs in the total TTF1-positive cell population was measured for each of the 50 lesions in PDF and Agglomeration, and shown as the Tm4sf1 positive index in AEC2 (mean ± S.D.).

Further examination revealed that PU.1 (nucleus) and CD68 (cytoplasm) double positive macrophages were present in the PDF (Fig. 9A). In addition to AMs (Green frame areas in Fig. 9A, upper right, show an “Over-stuffed AM” in the alveolar air space), interstitial macrophage infiltration was observed in the alveolar interstitium within the PDF (Fig. 9A, upper right), and particle-laden interstitial macrophages (Blue frame areas in Fig. 9A, upper right) were scattered in the alveolar interstitium.

The number of CD34-positive vascular endothelial cells was severely decreased in the alveolar interstitium within the PDF compared to the surrounding normal alveolar interstitium (Fig. 9A, lower left). An increase of vascular endothelial growth factor receptor 3 (VEGFR3)-positive lymphatic vessels (Fig. 9A lower panels) was also not observed in the PDF. In addition, α-smooth muscle actin (αSMA)-positive myofibroblasts (Fig. 9A, lower right) and collagen fibers (Fig. S9) were also not found in the PDF.

Transmembrane 4 superfamily member 1 (Tm4sf1)-positive AEC2 (Alveolar Epithelial Progenitor, AEP) appears during tissue regeneration of lung injury [34, 35]. In the present study, we determined the AEP index in the “Agglomeration” and “PDF” lesions to ascertain whether AEP regeneration of lung injury was distinct in these two lesions. We found that the AEP positive index was the same in these lesions (Fig. 9B) and dramatically higher than in the surrounding normal alveolar region (Fig. S8). These results suggest that Agglomeration is an initial change leading to the development of PDF, and consequently, that the AEC2 to AEP transformation and epithelial proliferation in the PDF appear to be the result of a reaction to the particle-laden macrophages that constitute the Agglomeration lesion.

Finally, the incidence and the multiplicity of PDF were examined. The incidence was significant in males exposed to 25 and 50 mg/m^3^ and in females exposed to 12.5, 25, and 50 mg/m^3^ (Table 2). Incidence and multiplicity showed an exposure concentration-dependent increase in both sexes. In addition, in the 50 mg/m^3^ exposure groups, there was a statistically significant increase in the multiplicity of PDF in females compared to males (Table 2), indicating that there is a sex difference in the development of PDF.

**Table 2.** Incidence and multiplicities of the pulmonary dust foci.

| Concentration<br>(mg/m<sup>3</sup>) | No. of<br>rats | Pulmonary dust foci |  |  |  |  |  |
| --- | --- | --- | --- | --- | --- | --- | --- |
|  |  | Incidence<br>(%) | Multiplicities (average ± SD) |  |  |  |  |
|  |  |  | No./slide |  | No./cm<sup>2</sup> lung |  |  |
| Male |  |  |  |  |  |  |  |
| 0 | 10 | 0 (0) | 0 |  | 0 |  |  |
| 6.3 | 10 | 0 (0) | 0 |  | 0 |  |  |
| 12.5 | 10 | 1 (10) | 0.10 ± | 0.32 | 0.07 ± | 0.21 |  |
| 25 | 10 | 5 (50)<sup>\$\$</sup> | 2.10 ± | 2.60 | 1.12 ± | 1.36 | |
| 50 | 10 | 10 (100)<sup>\$\$\$</sup> | 76.40 ± | 18.37<sup>***</sup> | 47.85 ± | 15.65<sup>***</sup> | |
| Female |  |  |  |  |  |  |  |
| 0 | 10 | 0 (0) | 0 |  | 0 |  |  |
| 6.3 | 10 | 2 (20) | 0.20 ± | 0.42 | 0.17 ± | 0.36 |  |
| 12.5 | 10 | 4 (40)<sup>\$</sup> | 0.40 ± | 0.52 | 0.33 ± | 0.42 | |
| 25 | 10 | 9 (90)<sup>\$\$\$</sup> | 7.80 ± | 4.44 | 5.37 ± | 3.23 | |
| 50 | 10 | 10 (100)<sup>\$\$\$</sup> | 99.00 ± | 25.43<sup>***,###</sup> | 70.89 ± | 21.03<sup>***,###</sup> | |

Significant difference: \$, p<0.05; \$\$, p<0.01; \$\$\$, p<0.001 by Chi square test compared with each cont.
\*\*\*, p<0.001 compared with each cont or ###, p<0.001 compared with male 50 mg/m<sup>3</sup> by tukey's multiple comparison test .

### Cell proliferation ability in AEC2 in PDF

As described in the previous section, in the PDF lesion AEC2 hyperplasia is co-localized with an agglomeration of particle-laden macrophages. Since PDF is a major lesion caused by TiO_2_ NP inhalation, it is important to know the cell proliferative activity of the AEC2 cells within the PDF. We performed double staining for Ki67, a cell proliferation marker, and LPCAT1, an AEC2 marker, to evaluate the proliferation activity of AEC2 (Fig. 10). The results showed that the AEC2 Ki67-positive index in the PDF of both sexes in the 50 mg/m^3^ groups is significantly higher compared to both the alveolar area of rats in the clean air group (0 mg/m^3^) and in the tissue surrounding a lesion (SUR) (Fig. 10B). In addition, the Ki67-positive index within the PDF was significantly higher in females than in males. These results indicate that, in the PDF induced by TiO_2_ NP, AEC2 has increased cell proliferative activity and that there is a sex difference in AEC2 proliferation.

**Figure 10.**
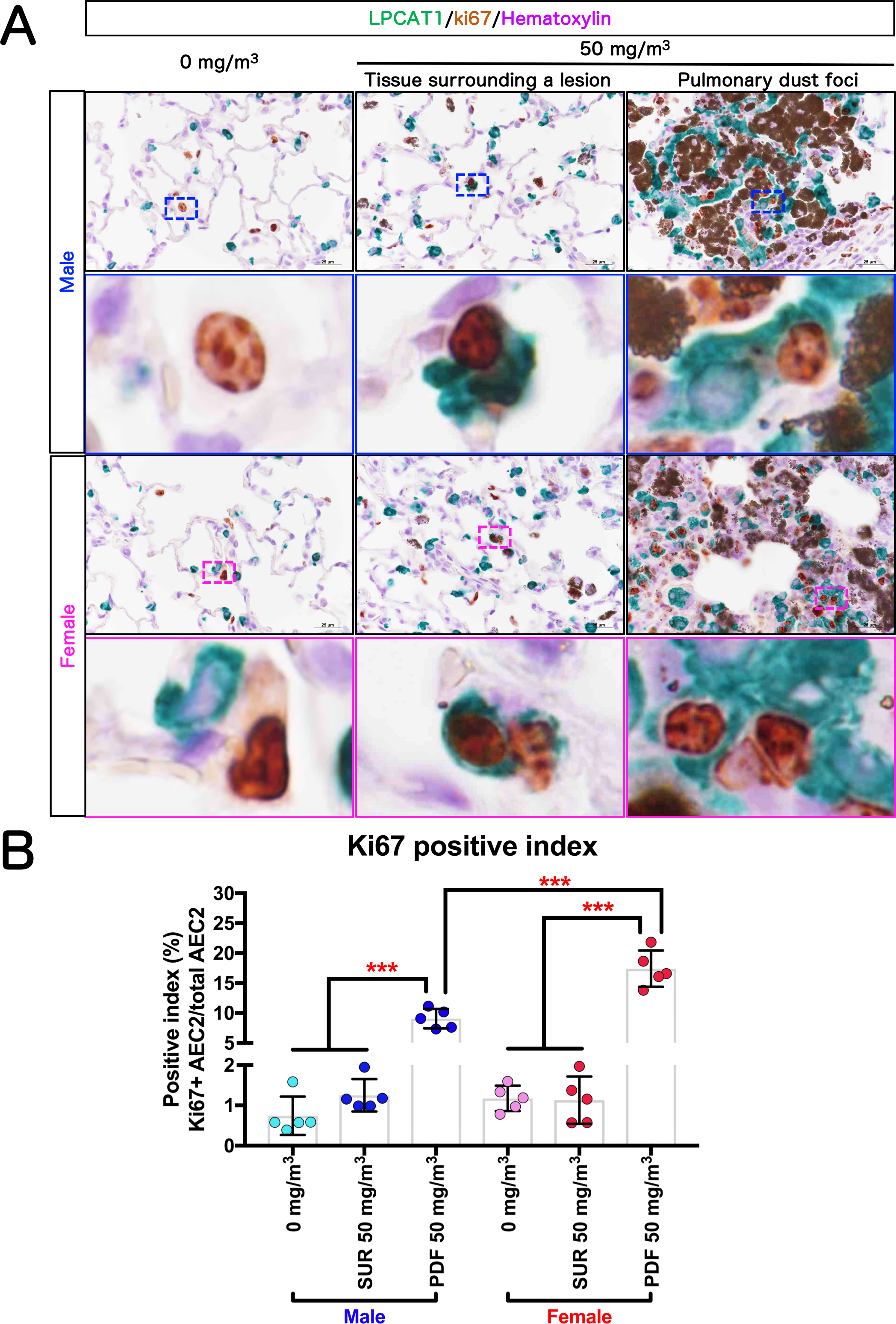
Cell proliferative activity of AEC2. Representative immunohistochemical staining images of Ki67, a cell proliferation marker, and LPCAT1, an AEC2 marker (A). The Ki67-positive index in AEC2 was calculated as the percentage of Ki67 and LPCAT1 double positive cells compared to the total LPCAT1-positive cell population (B). Statistical significance was analyzed using Tukey’s multiple comparison test: \*\*\**p*<0.001 indicate significant differences. Abbreviations: PDF: pulmonary dust foci and SUR: tissue surrounding a lesion.

### DNA damage in AEC2 in PDF

The postulated AOP of TiO_2_-induced pulmonary toxicity suggests that persistent inflammation (KE3) leads to epithelial cell proliferation (KE6) via injury to lung epithelial cells (KE4) and that proliferation of epithelial cells (KE6) with DNA damage (KE5) leads to preneoplastic epithelial lesions (KE7) [31]. Therefore, we assessed DNA damage in AEC2 by double staining for phosphorylation of the Ser-139 residue of the histone variant H2AX (γ-H2AX), a DNA double-strand break marker, and LPCAT1, an γ AEC2 marker (Fig. 11). Consistent with the results in Fig. 10 showing AEC2 proliferation (KE6), the γ-H2AX-positive index of the AEC2 in the PDF was significantly increased in both sexes in the 50 mg/m^3^ groups compared to both the alveolar area of rats in the clean air group (0 mg/m^3^) and in the tissue surrounding a lesion (SUR) (Fig. 11B), and the γ-H2AX-positive index was significantly higher in females than in males. These results indicate that AEC2 in PDFs acquire DNA damage and that a higher proportion of cells are damaged in females.

**Figure 11.**
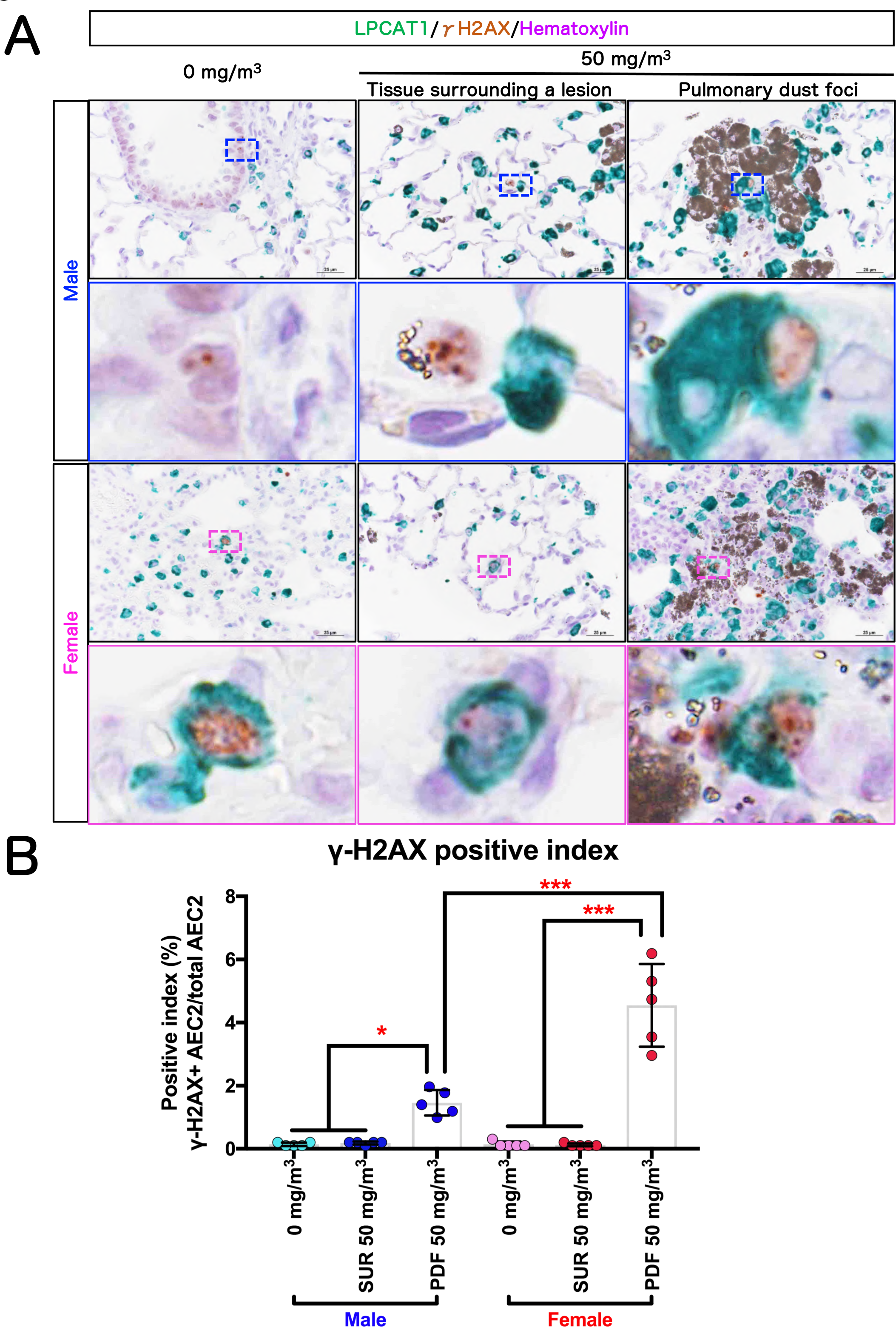
DNA damage in AEC2. Representative immunohistochemical staining images of the Ser-139 residue of the histone variant H2AX (γ-H2AX), a DNA double-strand break marker, and LPCAT1, an AEC2 marker (A). The γ-H2AX -positive index in AEC2 was calculated as the percentage of γ-H2AX and LPCAT1 double positive cells compared to the total LPCAT1-positive cell population (B). Statistical significance was analyzed using Tukey’s multiple comparison test: \**p*<0.05 and \*\*\**p*<0.001 indicate significant differences.

### Histopathological findings in other organs

Histopathological findings in the nasal cavity, nasopharynx, heart, liver, kidney, pituitary gland, thyroid, testis, epididymis, prostate, oviduct, eye, Harderian gland, and bone marrow are shown in Table S5. Inhalation exposure to TiO_2_ NPs caused toxic changes in the nasal cavity and nasopharynx. In the nasal cavity and nasopharynx, goblet cell hyperplasia was observed in male rats exposed to 25 and 50 mg/m^3^ TiO_2_ NP and in female rats exposed to 12.5, 25, and 50 mg/m^3^ TiO_2_ NP. Eosinophilic changes in the olfactory and respiratory epithelium were induced in both sexes exposed to 12.5, 25, and 50 mg/m^3^ TiO_2_ NP. Deposition of particles in the lymphoid tissue was observed in all exposed groups.

## Discussion

We conducted a 13-week inhalation toxicity study of anatase TiO_2_ NPs according to OECD TG 413 guidelines. One of the goals of the study was to identify and assess rat pulmonary lesions associated with pneumoconiosis. Male and female rats were exposed to 6.3, 12.5, 25 and 50 mg/m^3^ TiO_2_ NP for 6 hours per day, 5 days per week, for 13 weeks. Evaluation of all organs in the rats demonstrated that damage caused by inhalation exposure to TiO_2_ NP was limited to the respiratory tract, especially the lung. The TiO_2_ NPs exposed lungs showed multiple milky white spots on gross examination. Histopathological examination identified the main lesions as an agglomeration of particle-laden macrophages and AEC2 hyperplasia. We defined these lesion as pulmonary dust foci (PDF). As discussed below, PDF are likely to be early lesions associated with pneumoconiosis (Fig. 12).

**Figure 12.**
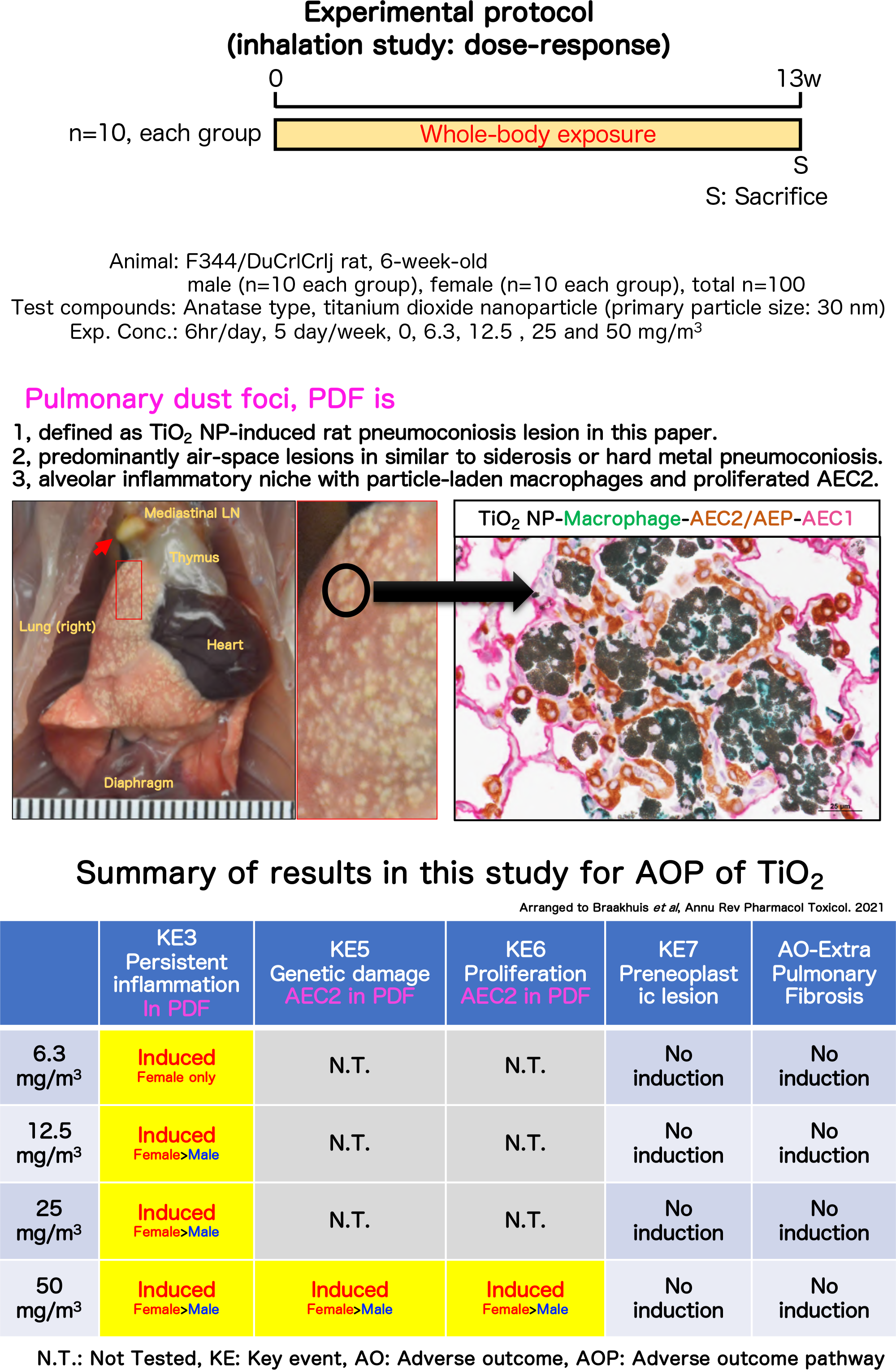
Graphical abstract in this study.

The PDF has characteristics of an inflammatory niche with neutrophil and lymphocyte infiltration, decreased vascular endothelium, and particle-laden interstitial macrophages infiltration in the interstitium. The AEC2 in the PDF express Tm4sf1, indicating that they are Alveolar Epithelial Progenitor cells (AEP) and this type of cell contributes to alveolar regeneration. In agreement with this data, the AEP in the PDF had an increased Ki67-positive index indicating proliferative activity. PDF were observed in males exposed to 12.5 mg/m^3^ and higher concentrations of TiO_2_ NP and in females exposed to 6.3 mg/m^3^ and higher concentration of TiO_2_ NP. PDF developed in both sexes in an exposure concentration-dependent manner. The finding of PDF is a key event in the pulmonary toxicity due to inhalation exposure to TiO_2_, and can be regarded as persistent inflammation. Therefore, that the lowest observed adverse effect concentration (LOAEC) for TiO_2_ NPs in this study was 12.5 mg/m^3^ for males and 6.3 mg/m^3^ for females. The no observed adverse effect concentration (NOAEC) was 6.3 mg/m^3^ for males. In females, the lowest exposure concentration of TiO_2_ NPs caused the formation of PDF, therefore, in this study a NOAEC in females was not found.

The present study demonstrated sex differences in TiO_2_ NP-induced lung toxicity at the highest dose of 50 mg/m^3^: increases in LDH activity, total protein and albumin levels in the BALF, the incidence and multiplicity of Ki67- and γ-H2AX-positive indices in the PDF were higher in females than in males exposed to 50 mg/m^3^ TiO_2_ NP. Notably, the lung burden tended to be higher in females than males. These results suggest that there is a sex difference in the onset and progression of TiO_2_ toxicity, with females being more susceptible to TiO_2_ toxicity than males, seemingly in part due to differences in retention of TiO_2_ due to impaired clearance. The fact that several toxicity indicators, including LDH activity, correlated positively with lung burden support this possibility.

It is known from previous epidemiology and case reports of workers that pneumoconiosis can be caused by inhalation of various materials, including asbestos [36], silica [37], mixed dust [38], hard metals [39, 40], aluminum [41], beryllium [42], indium [43–45], and talcum [46]. In addition to the clinical findings of pneumoconiosis due to inhalation of the materials listed above, there have been many case reports of workers exposed to TiO_2_ particles including nanoparticles and grindings [11–22]. These TiO_2_ associated lesions in workers’ lungs ranged from alveolar lesions with agglomeration of particle-laden macrophages both in the air space and the alveolar interstitium, granulomas, alveolar fibrotic lesions such as nonspecific interstitial pneumonia (NSIP), bronchitis, and alveolar proteinosis. The variety of different lesions caused exposure to TiO_2_ particles is likely due to the type of TiO_2_ particles the workers are exposed to. In addition, various confounding factors such as contamination of TiO_2_ particles with other minerals and smoking history, contribute to mixed reactions to inhaled TiO_2_ [23]. However, histopathological comparisons of rat lung lesions obtained in this study and pneumoconiosis of workers exposed to TiO_2_, it is evident that several of these workers have alveolar lesions similar to rat PDF [18]. In addition, many workers’ pneumoconiosis, such as arc-welders lung (also known as pulmonary siderosis) [47–50] and hard metal pneumoconiosis [39,40,51], have also been reported to have histopathological characteristics similar to rat PDF composed both of hypertrophic alveolar epithelial proliferation and alveolar filling macrophages. Furthermore, there are many clinical findings of alveolar lesions with PDF-like histopathology in idiopathic pulmonary hemosiderosis [52] and smoking-related lesions, such as smoking-related interstitial fibrosis (SRIP) [53, 54] and respiratory bronchiolitis interstitial lung disease/desquamative interstitial pneumonia (RBILD/DIP) [55–58]. Additionally, similar alveolar lesions occur in experimental animals such as rats and rabbits exposed by inhalation to different types of TiO_2_ [29, 59]. In summary, the PDF observed in this rat study is an early lesion of pneumoconiosis caused by exposure to TiO_2_, and is likely to be an alveolar reaction common to mammals. Further observation is necessary to determine whether PDF progresses to fibrotic interstitial lung disease over time.

In this study, we also examined whether the presumed key elements of the AOP caused by TiO_2_ inhalation occurred within the PDF (Fig. 12). For this analysis, we used lung samples from the male and female 50 mg/m^3^ exposure groups. The results showed inflammatory cells localized in the PDF, indicating that the PDF is an inflammatory niche where inflammatory cells infiltrate with particle-laden macrophages, and is a “microenvironment” where persistent inflammation (KE3) occurs. A significant increase in γ-H2AX and Ki67 positive indices in AEC2 in the PDF but not the surrounding area of the lesions provides clear evidence of genetic damage of lung epithelial tissue (KE5) and AEC2 proliferation (KE6) in the PDF lesions (Fig. 12). Our findings strongly support a mechanism whereby particle-laden macrophages become lodged in the alveolar airspace causing, which leads to persistent inflammation (KE3), persistent epithelial injury (KE4), and regenerative proliferation of AEC2 (KE6). Although in the present study, γ-H2AX expression was increased specifically in AEC2 in the PDF of the 50 mg/m^3^ exposure group, previous reports measuring TiO_2_ inhalation mediated DNA damage have all been negative [31,60,61], and the intratracheal instillation studies of TiO_2_ have not been consistent [31,62–67]. Therefore, the present study is the first report not only to define and discuss the histopathological and cell biological basis of TiO_2_ NPs-induced rat lung lesions caused by inhalation exposure to TiO_2_ NPs but also to demonstrate clearly increased DNA damage in AEC2. In the present study neither typical preneoplastic lesions (KE7) nor pulmonary fibrotic lesions (AO-Extra) were observed in either the male or female 50 mg/m^3^ exposure groups. As mentioned above, the PDF can be considered to be the initial lesion of TiO_2_ NP-induced pneumoconiosis in rats. Since the progression of pneumoconiosis in workers is well known to be complicated by lung cancer [27, 28], it is important to investigate whether lung cancer in rats will develop as a complication of the PDF. To address this issue, we are currently conducting a 104-week systemic inhalation study using F344 rats.

## Conclusions

Inhalation exposure to TiO_2_ NPs for 13 weeks induced pulmonary lesions triggered by particle-laden macrophages in the alveoli of the F344 rat lung. We defined this specific lesion as pulmonary dust foci (PDF). The TiO_2_ NP-induced rat PDF is an inflammatory niche in the lung. Persistent inflammation causes tissue damage and induces AEC2 transformation to alveolar epithelial progenitor cells (AEP) which proliferate to repair inflammation mediated tissue damage. In the presence of inflammatory mediators AEP cells acquire DNA damage. Based on PDF induction, the LOAEC for pulmonary disorders in male and female rats in this study was 12.5 mg/m^3^ and 6.3 mg/m^3^, respectively. There was a sex difference in the lung lesions onset, with females showing more progressive lesion parameters than males. The similar histopathology to human pneumoconiosis makes it is highly likely that the PDF in the rat is an early lesion of rat pneumoconiosis. Further studies should focus on the progression of PDF over time for better understanding of TiO_2_-NP-inhalation-mediated pneumoconiosis due to and its carcinogenic potential. Importantly, different TiO_2_ particles will have different toxicities. For example, LDH activity in rat BALF after inhalation exposure to TiO_2_ NP (this study) was lower than that of ultrafine TiO_2_ particles reported in a previous study [29] (Table S6), suggesting that inhalation of the anatase type TiO_2_ NP using in this study is less harmful than inhalation of P25.

## Materials

Anatase type nano-titanium dioxide, TiO_2_ NP (aNTiO_2_) (Fig. S10) was purchased from Tayca co. (primary particle size: 30 nm). A list of all primary antibodies used in these studies is summarized in Table S7. Other reagents used in the study were of the highest grade available commercially.

### Animals

Male and female F344 rats at 4 weeks old were purchased from Charles River Laboratories Japan, Inc. (Kanagawa, Japan). The rats were housed in an air-conditioned room under a 12 hours light/12 hours dark (8:00-20:00, light cycle) photoperiod, and fed a general diet (CR-LPF, Oriental Yeast Co. Ltd., Tokyo, Japan) and tap water *ad libitum*. After a 1 week quarantine and acclimation period, they were exposed to TiO_2_ NP. All animal experiments were approved by the Animal Experiment Committee of the Japan Bioassay Research Center.

### Generation of TiO_2_ NP aerosol

The generation of TiO_2_ NP aerosol into the inhalation chamber was performed using our established method (cyclone sieve method) [68, 69] with some modifications. Briefly, TiO_2_ NP was fed into a dust feeder (DF-3, Shibata Scientific Technology, Ltd., Soka, Japan) to generate TiO_2_ NP aerosol, and the aerosol was introduced into a particle generator (custom-made by Seishin Enterprise Co., Ltd., Saitama, Japan) to separate the aerosol and feed it into the inhalation chamber. The concentration of the TiO_2_ NP aerosol in the chamber was measured and monitored by an optical particle controller (OPC; OPC-AP-600, Shibata Scientific Technology), and the operation of the dust feeder was adjusted by feedback control based on upper and lower limit signals to maintain a steady state.

The mass concentration of TiO_2_ NP aerosol in the chamber was measured every two weeks during the exposure period. Aerosols collected on a fluoropolymer binder glass fiber filter (T60A20, φ55 mm, Tokyo Dylec, Corp., Tokyo, Japan) were weighed for each target concentration at 1, 3, and 5 hours after the start of the exposure. Using the mass per particle (K-value) calculated using the measured mass results (mg/m^3^) and the particle concentration data (particles/m^3^) obtained from the OPC, the particle concentration for each group during the exposure period was converted to a mass concentration. The particle size distribution and morphology of the TiO_2_ NPs were measured at the 1st, 13th, and 25th weeks of the exposure. The particle size distribution was measured using a micro-orifice uniform deposit cascade impactor (MOUDI-II, MSP Corp., Shoreview, MN). The MMAD and σg were calculated by cumulative frequency distribution graphs with logarithmic probability (Fig. S1C). The TiO_2_ NP in the inhalation chamber was collected on a 0.2 μm polycarbonate filter ( 47 mm, φ Whatman plc, Little Chalfont, UK), and observed using SEM (SU8000, Hitachi High-Tech, Tokyo, Japan).

### 13-week inhalation study

This experiment was conducted with reference to the OECD Guideline for Testing of Chemicals (TG 413) [70]. Based on the results of a dose-finding study conducted previously and the OECD TG 413, target concentrations for TiO_2_ NP aerosols were set at 6.3, 12.5, 25, and 50 mg/m^3^, and the exposure schedule was 6 hours per day, 5 days per week, for 13 weeks (Fig. S11). One hundred rats (10 males and 10 females) in each group were transferred to individual stainless steel cages and exposed to TiO_2_ NP for 6 hours with access to food or water. At 1-2 days after the last exposure, the rats were fasted for 1 day and the blood of the rats was collected under isoflurane anesthesia, and the rats were euthanized by exsanguination. BALF was collected from five males and five females in each group as described below. For histopathological analysis, all tissues were collected from all the rats in each group, and fixed in 10% neutral phosphate buffered formalin solution.

### BALF collection and analysis

The left bronchus was tied with a thread, and the right lung was lavaged: 4∼5ml of saline was injected into the lung through the trachea, in and out twice, and collected as BALF. The total cell numbers in the BALF were counted using an automatic cell analyzer (ADVIA120, Siemens Healthcare Diagnostics Inc. Tarrytown, NY). Thin-layer cell populations on glass slides were prepared using Cytospin 4 (Thermo Fisher Scientific, Inc., Waltham, MA). After May-Grunwald-Giemsa staining, differential white blood cell count was made by visual observation. BALF cytospin specimens were carefully examined under a microscope to classify the status of AMs phagocytosing TiO_2_ NPs. All AMs were divided into TiO_2_ NPs-laden AMs and normal AMs. The TiO_2_ NPs-laden AMs were then classified as Over-stuffed AMs, which had phagocytosed TiO_2_ NPs until the nucleus was no longer visible, Burst AMs, which were disintegrated into particles and cellular debris, and the number of each AM was counted.

BALF was centrifuged at 1,960 rpm (800 × *g*) for 10 min at 4°C, and the activity of LDH, ALP and γ-GTP, and the level of total protein, albumin in the supernatant was measured using an automatic analyzer (Hitachi 7080, Hitachi, High-Tech Corp., Tokyo, Japan).

### Titanium burden analysis

To determine the lung burden of Ti in TiO_2_ NP-exposed rats, the lung pieces was collected and weighed. The lung was put into glass vessels, treated with 3 mL of distilled water 3 mL of sulfuric acid and 1 mL of nitric acid at 270°C for 1 h. Samples were then diluted to 30-50 mL with 3% sulfuric acid. The samples were further diluted 2 to 50 fold to keep the concentration within the calibration curve, and TiO_2_ concentration in the samples was determined by a Zeeman atomic absorption spectrometry (Z-5010; Hitachi High-Tech Corporation, Tokyo, Japan) with a Hitachi High-Tech lamp for Ti (part#207-2012 Serial 0490158100). Absorbance of the digested samples was detected at 364.3 nm. Quantification was performed using a seven point calibration curve prepared by diluting appropriate volumes of a 1000 mg/L stock solution (Kanto Chemical Co., Inc., Tokyo, Japan) for 0.025, 0.05, 0.1, 0.15, 0.2, 0.3 and 0.4 µg/ml. TiO_2_ concentrations were calculated from the corresponding molecular weight ratio of TiO_2_ to Ti. The values obtained were calculated as the amount of Ti per gram. The Pearson correlation coefficient (Pearson’s r) was calculated between the lung burden and several toxicological markers using GraphPad Prism 5 (GraphPad Software, San Diego, CA).

### Hematological and blood chemistry tests

For hematological examination, blood samples collected at the time of each autopsy were analyzed with an automated hematology analyzer (ADVIA120, Siemens Healthcare Diagnostics Inc. Tarrytown, NY). For biochemical tests, the blood was centrifuged at 3,000 rpm (2,110 × *g*) for 20 minutes, and the supernatant was analyzed with an automated analyzer (Hitachi 7080, Hitachi, Ltd., Tokyo, Japan).

### Histopathological analysis

Serial tissue sections were cut from paraffin-embedded lung specimens, and the first section (2-μm thick) was stained with H&E for histological examination and the remaining sections were used for immunohistochemcal analysis. The histopathological finding terms used in this study for lesion were determined by certified pathologists from the Japanese Society of Toxicologic Pathology, based on the terms adopted by International Harmonization of Nomenclature and Diagnostic Criteria for Lesions in Rats and Mice (INHAND)[71]. Pathological diagnosis was performed blindly by three pathologists and summarized as a result of the discussion. Each non-neoplastic lesion was evaluated for its severity and scored on a scale of “slight” to “severe” with reference to the criteria by Shackelford et al [72].

### Masson’s Trichrome staining

Details of this procedure have been described previously [34]. Briefly, the slides were deparaffinized, washed with water, and then reacted with an equal volume mixture of 10% potassium dichromate and 10% trichloroacetic acid for 60 minutes at room temperature. The specimens were then washed with water and stained with Weigelt’s iron hematoxylin solution (C.I.75290, Merck-Millipore) for 10 minutes at room temperature. Then successively stained with 0.8% orange G solution (C.I.16230, Merck-Millipore) reacted for 10 minutes at room temperature, Ponceau (C.I.14700, FUJIFILM-Wako Pure Chemical Corp., Osaka, Japan), acid fuchsin (C.I.42685, Merck-Millipore), and azofloxine (C.I.18050, Chroma Germany GmbH, Augsburg, Germany) mixture reacted for 40 minutes at room temperature, 2.5% phosphotungstic acid reacted for 10 minutes at room temperature, and blue aniline solution (C.I.42755, Chroma Germany GmbH) under a microscope. Between each staining solution the slides were washed lightly with 1% acetic acid in water. Then, dehydration, permeabilization, and sealing were performed.

### Elastica Van Gieson staining

Briefly, the slides were deparaffinized, washed with water, reacted with Maeda Modified Resorcinol-Fuchsin Staining Solution (Mutoh Chemical, Part No. 40321, Japan) for 30 min at room temperature, and separated with 100% ethanol. After washing with water, the slides were stained with Weigelt’s iron hematoxylin solution (C.I.75290, Merck-Millipore, US) for 10 minutes at room temperature and washed with running water for 10 minutes. The slides were then reacted with 1% Sirius red solution (Mutoh Chemical, Part No. 33061, Japan) for 3-5 minutes at room temperature, separated and dehydrated with 90%-100% ethanol, permeabilized, and sealed.

### Immunohistological multiple staining analyses

Details of the multiple staining method have been described previously [73]. Briefly, lung tissue sections were deparaffinized with xylene, hydrated through a graded ethanol series, and incubated with 0.3% hydrogen peroxide for 10 min to block endogenous peroxidase activity. Slides were then incubated with 10% normal serum at room temperature (RT) for 10 min to block background staining, and then incubated for 2 hr at RT with the first primary antibody. After washing with PBS, the slides were incubated with histofine simple stain ratMAX-PO(MULTI) (414191, Nichirei, Tokyo, Japan) for

30 min at RT. After washing with PBS, slides were incubated with DAB EqV Peroxidase Substrate Kit, ImmPACT (SK-4103, Vector laboratories) for 2-5 min at RT. Importantly, after washing with dH_2_O after color detection, the sections were treated with citrate buffer at 98°C for 30 min before incubation with the next primary antibody to denature the antibodies already bound to the section. This procedure was repeated for the second and then the third primary antibody. HighDef red IHC chromogen (ADI-950-142, Enzo Life Sciences, Inc., Farmingdale, NY) was used for the second coloration and Histogreen chromogen (AYS-E109, Cosmo Bio, Tokyo, Japan) for the third coloration. Coloration was followed by hematoxylin staining for 30-45 seconds. The slides were then processed for light microscopy. The sections were observed under an optical microscope ECLIPSE Ni (Nikon Corp., Tokyo, Japan) or BZ-X810 (Keyence, Osaka, Japan).

To perform various morphometric measurements on PDFs, only the 50 mg/m^3^ group of both sexes, which could ensure a sufficient number of PDF occurrences to be analyzed, were used in this study.

For measurement of Ki67 and γ-H2AX positive indices, the male and female 0 mg/m^3^ groups (n=5) and the 50 mg/m^3^ groups (n=5) were used for analysis. For the 50 mg/m^3^ exposure groups, positive indexes were counted separately for pulmonary dust foci (PDF) and tissue surrounding a lesion (SUR). In all animals, at least ten fields of view were measured using a 40x objective lens. More than 500 LPCAT1-positive AEC2 per individual were measured for Ki67 and 1000 LPCAT1-positive AEC2 (PDF: more than 500 AEC2) per individual were measured for γ-H2AX, and the mean value per individual was used for statistical analysis.

For Tm4sf1 positive index in PDF and Agglomeration, 50 lesions were randomly selected from each 50 mg/m^3^ exposure group of each sex, and the percentage of TTF1/Tm4sf1 double positive AEP and TTF1-single positive AEC2 were measured.

### Statistical analysis

Except in the case of incidence and integrity of histopathological lesions, the data comparisons among multiple groups were performed by one-way analysis of variance with a post-hoc test (Dunnett’s or Tukey’s multiple comparison test), using GraphPad Prism 5 (GraphPad Software). The incidences and integrity of lesions were analyzed by the chi-square test using GraphPad Prism 5 (GraphPad Software). All statistical significance was set at *p* < 0.05.

## Abbreviations

AEC1: alveolar epithelial type 1 cell
AEC2: alveolar epithelial type 2 cell
AEP: alveolar epithelial progenitor
ALP: alkaline phosphatase
AM: alveolar macrophage
AO: adverse outcome
AOP: adverse outcome pathway
αSMA: α-smooth muscle actin
BALF: bronchoalveolar lavage fluid
BALT: bronchus-associated lymphoid tissue
CCSP: club cell secretory protein
CGRP: calcitonin gene-related peptide
σg: geometric standard deviations
γ-GTP: γ-glutamyl transpeptidase
γ-H2AX: phosphorylation of the Ser-139 residue of the histone variant H2AX
HE: hematoxylin and eosin
IE: initiating event
INHAND: International Harmonization of Nomenclature and Diagnostic Criteria for Lesions in Rats and Mice
KE: key event
LDH: lactate dehydrogenase
LOAEC: lowest observed adverse effect concentration
LPCAT1: lysophosphatidylcholine acyltransferase 1
MMAD: mass median aerodynamic diameter
NOAEC: no observed adverse effect concentration
PDF: pulmonary dust foci
SEM: scanning electron microscope
Sox2: SRY-Box Transcription Factor 2
SUR: tissue surrounding a lesion
TiO_2_ NPs: titanium dioxide nanoparticles
Tm4sf1: transmembrane 4 superfamily member 1
TTF1: Thyroid Transcription Factor 1
VEGFR3: vascular endothelial growth factor receptor 3

## Declarations

Ethics approval and consent to participate

All animals were treated humanely and all procedures were performed in compliance with the Animal Experiment Committee of the Japan Bioassay Research Center.

## Consent for publication

All authors gave their consent for publication of this manuscript.

## Availability of data and materials

The datasets used and analyzed during the current study are available from the corresponding authors on reasonable request.

## Competing interests

The authors declare that they have no competing interests.

## Acknowledgments

We wish to thank Dr. David B. Alexander of Nanotoxicology project, Nagoya City University Graduate School of Medicine for his insightful comments and English editing.

## Funding

This research was financially supported by a grant-in-aid from the Japan Organization of Occupational Health and Safety.

## Author information

Japan Bioassay Research Center, Japan Organization of Occupational Health and Safety, Kanagawa, Japan

Shotaro Yamano, Yuko Goto, Tomoki Takeda, Shigeyuki Hirai, Yusuke Furukawa, Yoshinori Kikuchi, Tatsuya Kasai, Kyohei Misumi, Masaaki Suzuki, Kenji Takanobu, Hideki Senoh, Misae Saito, Hitomi Kondo, Yumi Umeda

## Contributions

S.Y. and Y.U. performed the experiments and analyzed the data. S.H., Y.K., T.K., K.M. and M.S. assisted with animal experiments. K.T., H.S., Y.U., and S.Y. performed histopathological diagnoses. M.S. and H.K. performed BALF sampling and dissection.

Y.G. T.T. and S.Y. analyzed and interpreted the data. S.Y. and Y.U. conceived, designed, and directed the study and interpreted the data. S.Y., Y.G., T.T., and Y.U. drafted and revised the manuscript. All authors approved the manuscript as submitted.

## Corresponding authors

Correspondence to Shotaro Yamano or Yumi Umeda

## Supplementary Information

**Additional file 1: Figure S1.**
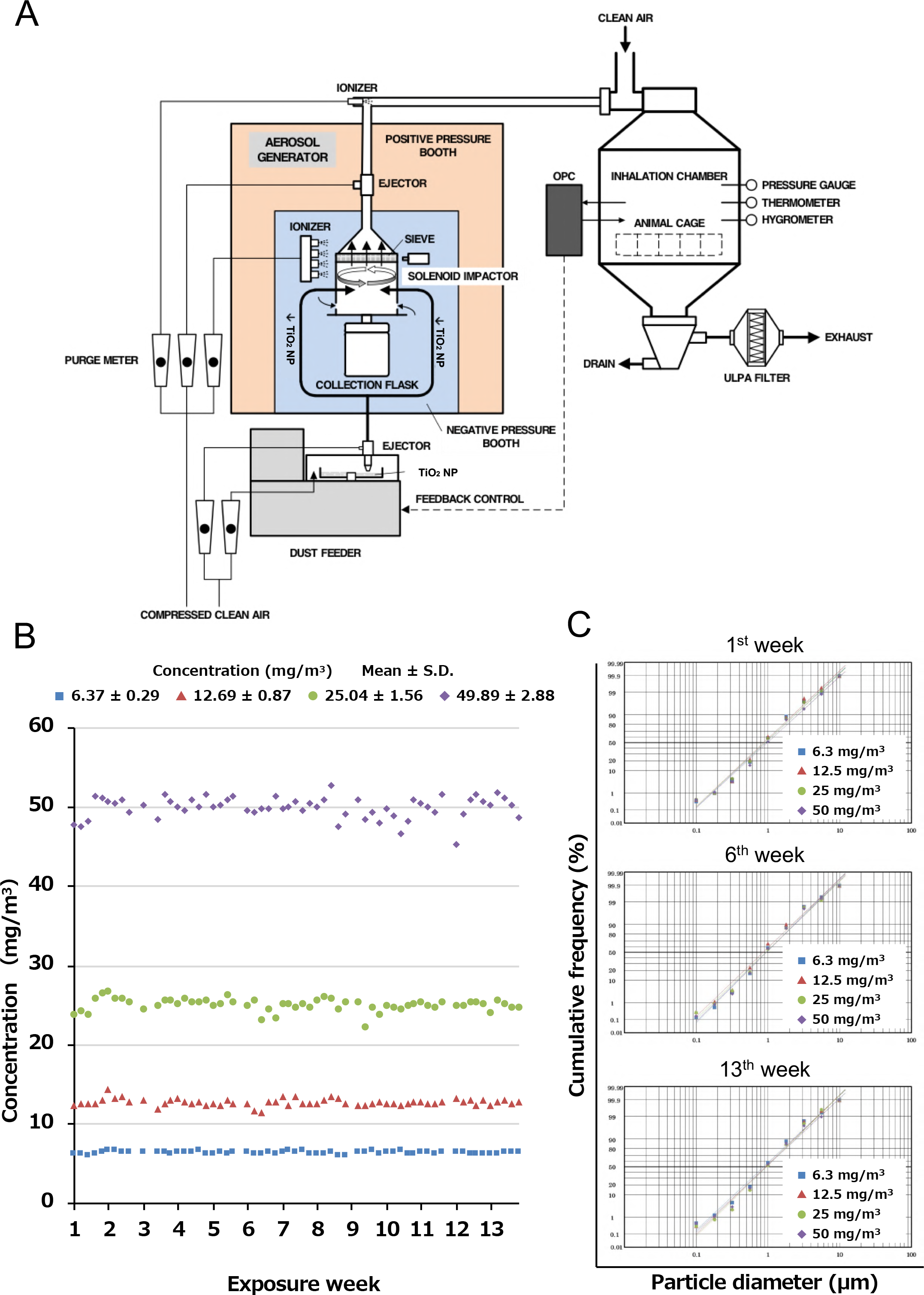
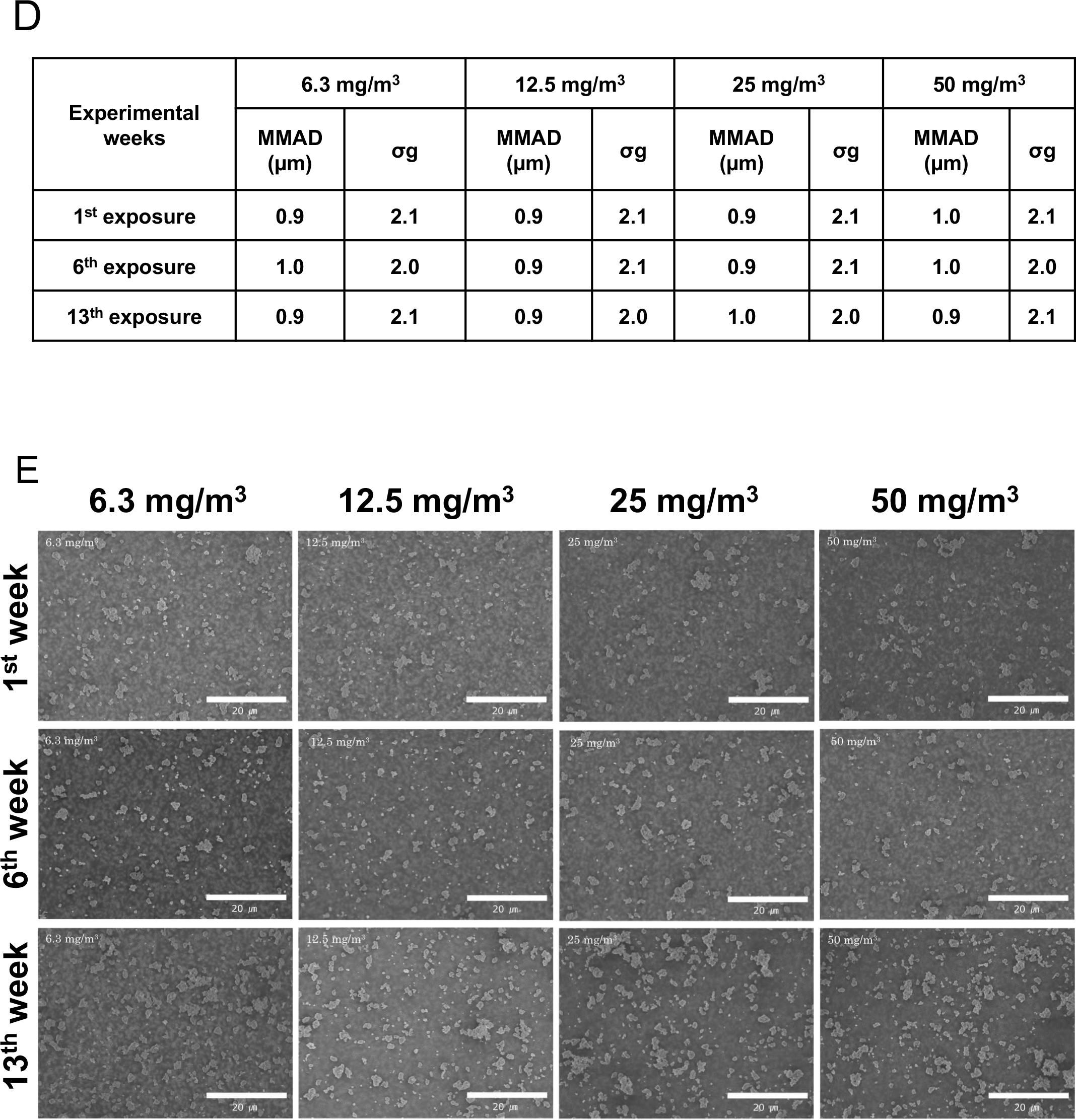
The whole body inhalation exposure system using in this study. The whole body inhalation exposure system (A). The averaged TiO_2_ NPs concentration in the chamber for each exposure day (B). Cumulative frequency distribution graphs with logarithmic probability (C). Mass median aerodynamic diameter (MMAD) and geometric standard deviation ( g) in the chamber (D). Representative scanning electron σ microscope (SEM) images of the TiO_2_ NPs in the chambers (E). Scale bar: 20 μm (panel E).

**Additional file 2: Figure S2.**
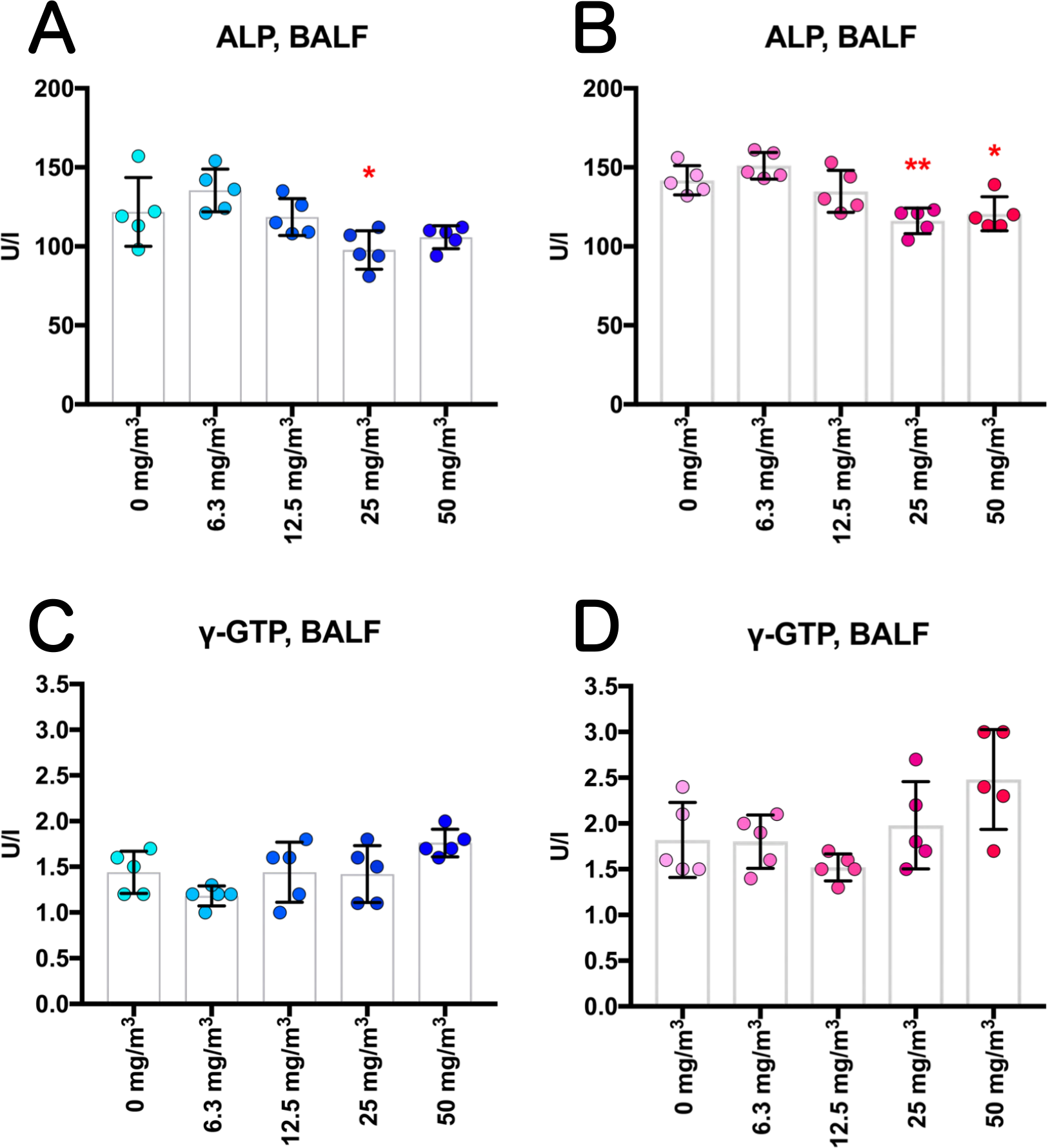
Biochemical markers in the BALF obtained from the lungs of rats after inhalation of TiO_2_ NPs for 13 weeks. Alkaline phosphatase (ALP) activity (A, B) and γ-Glutamyl transpeptidase (γ-GTP) activity (C, D) were measured using an automatic analyzer, and are shown by sex (males: A and C; females: B and D). Statistical significance was analyzed using Dunnett’s multiple comparison test: \**p*<0.05and \*\**p*<0.01.

**Additional file 3: Figure S3.**
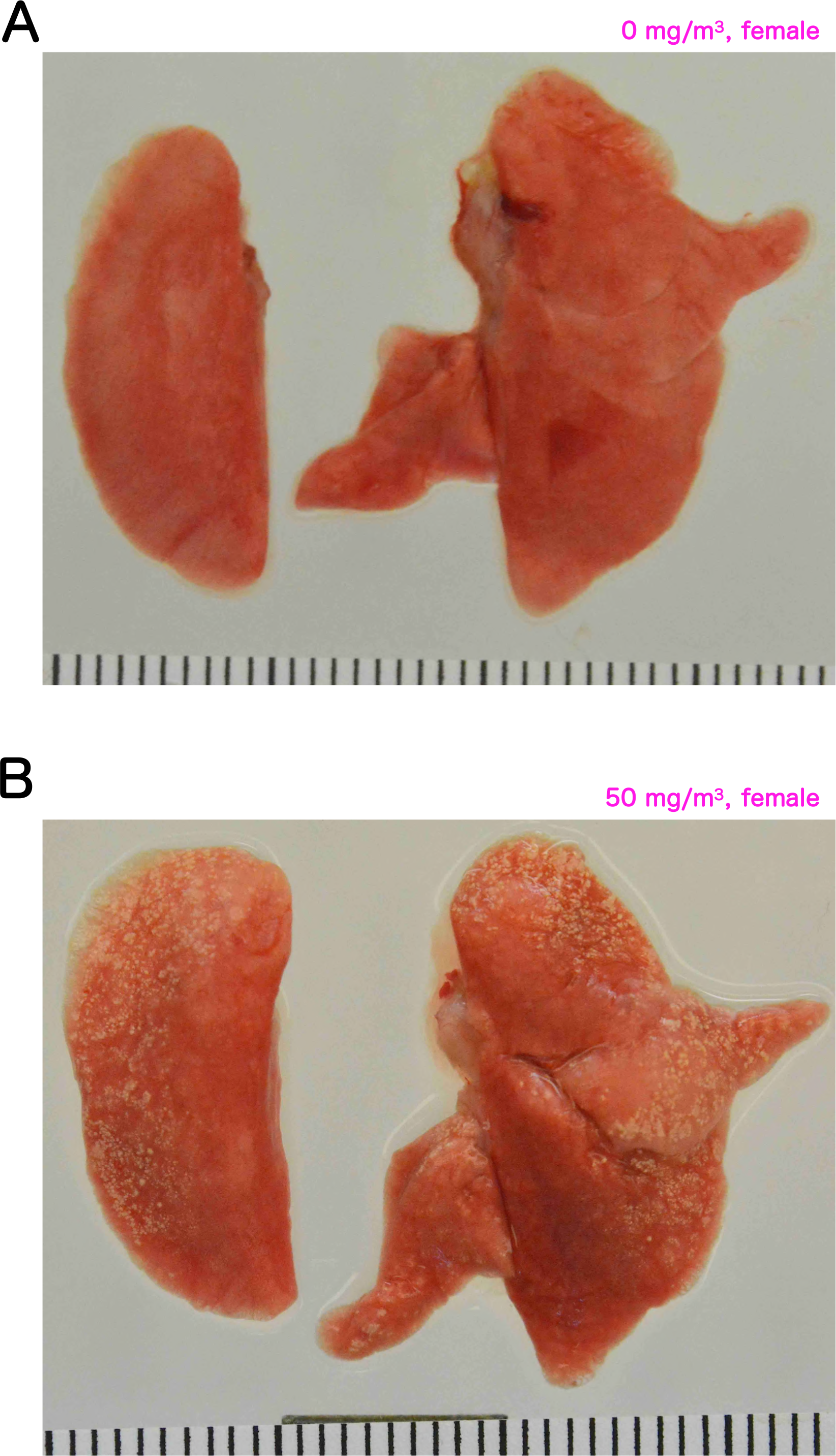
Representative macroscopic photographs of whole lungs. A: normal lungs of a female rat (0 mg/m^3^). B: TiO_2_ NPs-exposed lungs of a female rat (50 mg/m^3^)

**Additional file 4: Figure S4.**
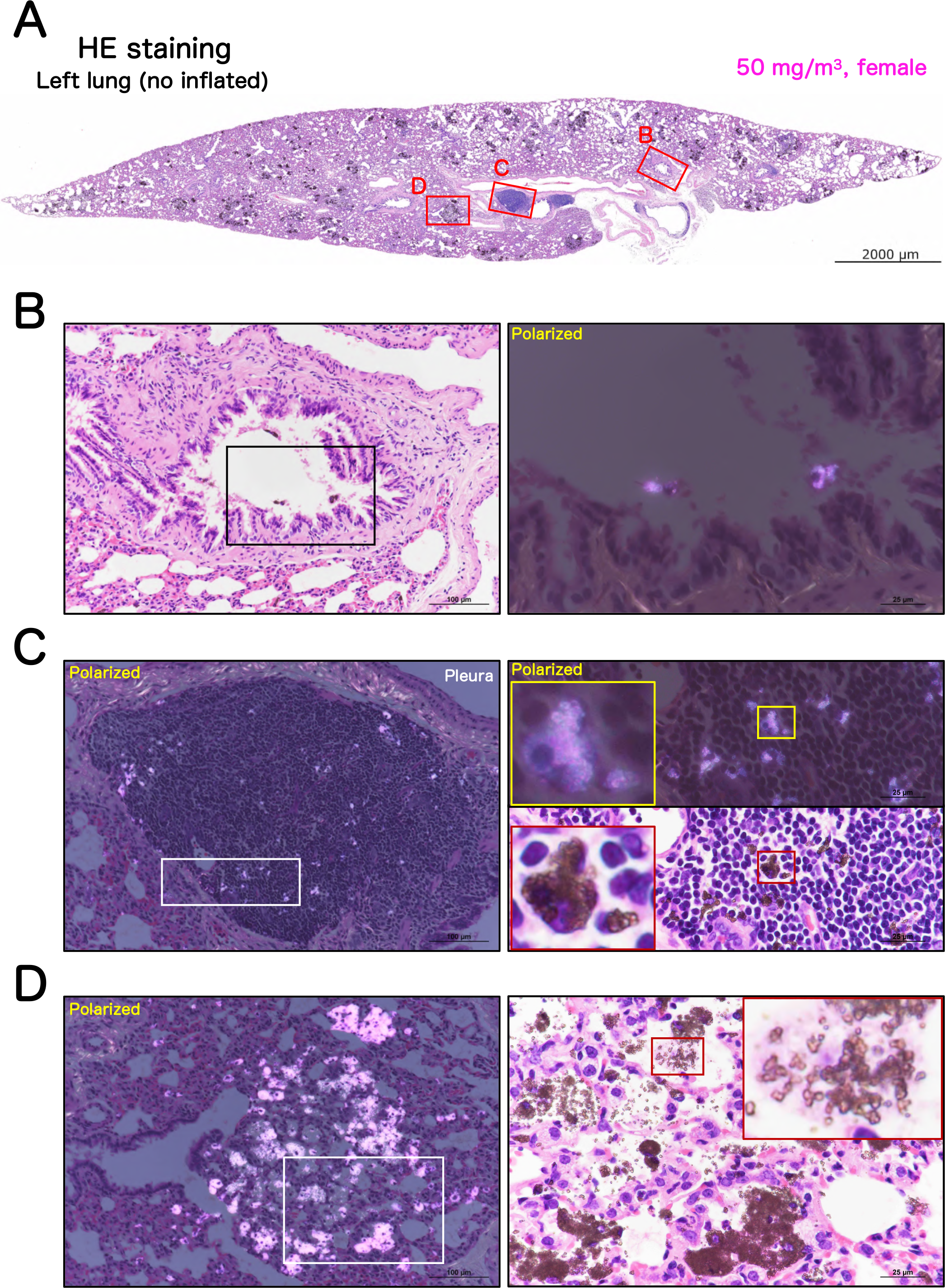
Representative microscopic photographs of a female rat left lung after inhalation exposure to TiO_2_ NPs (50 mg/m^3^): same rat as shown in figure 8. The left lung was not injected with formalin through the bronchus into the lung, and formalin immersion fixation was performed after the lung was removed. A typical loupe image (A) of the entire left lung and magnified images of each lesions (B-D). Particles in the process of being eliminated by the mucociliary escalator were observed on the bronchial mucosa (B). The infiltration of naked TiO_2_ NPs or particle-laden macrophages in bronchus-associated lymphoid tissue (BALT) (C). Burst macrophages were scattered in the 50 mg/m^3^ group of both sexes (D).

**Additional file 5: Figure S5.**
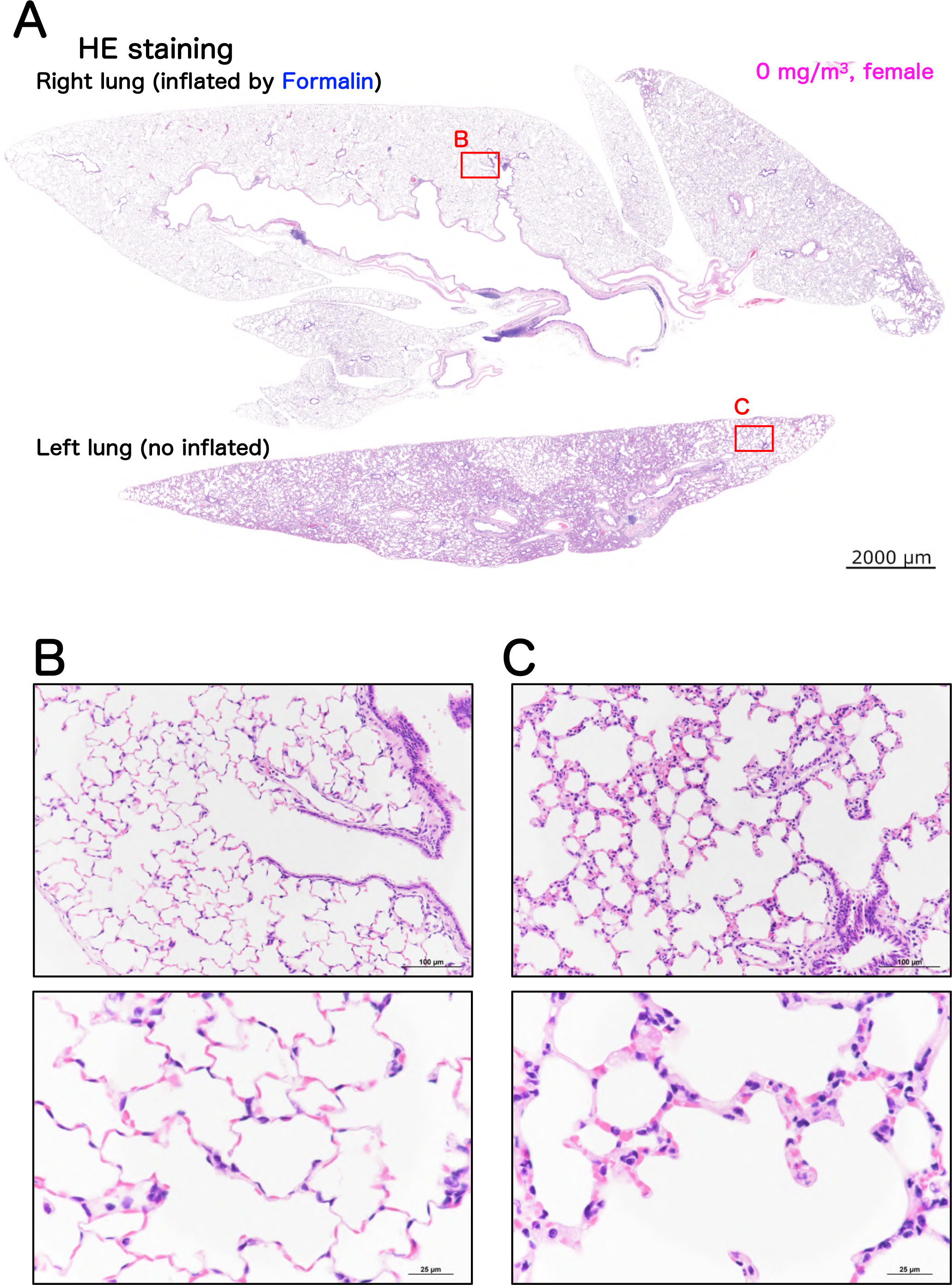
Representative microscopic photographs of the lungs of a female control rat. The lungs were stained with hematoxylin and eosin (HE) (see the Fig. 8 legend for details). A typical loupe image (A) of the entire lungs and magnified images of normal alveolar regions (B and C) are shown.

**Additional file 6: Figure S6.**
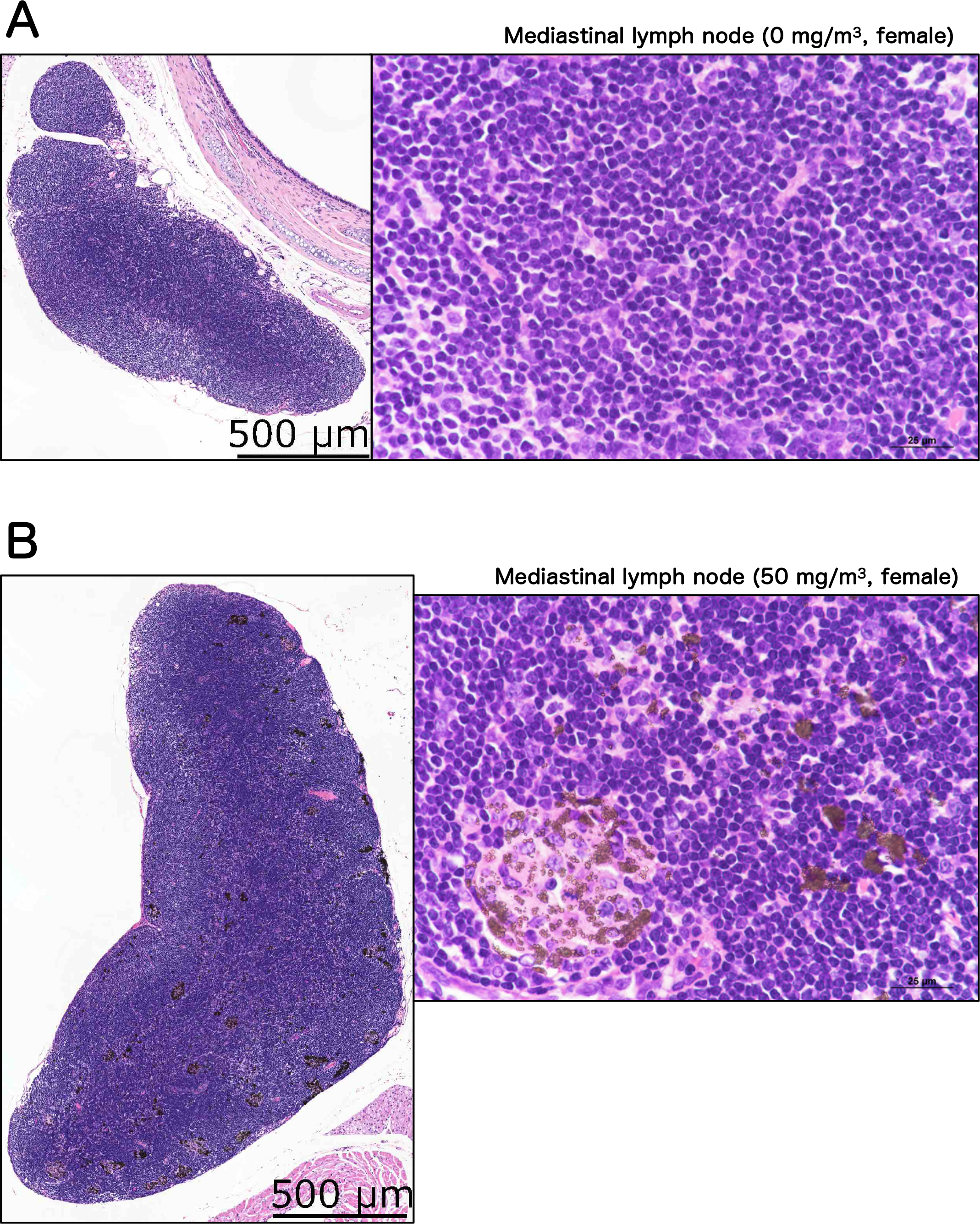
Representative microscopic photographs of mediastinal lymph node. A typical loupe image of the entire mediastinal lymph node and magnified images of each lymph node from a female control (A) and a female exposed to 50 mg/m^3^ (B) are shown.

**Additional file 7: Figure S7.**
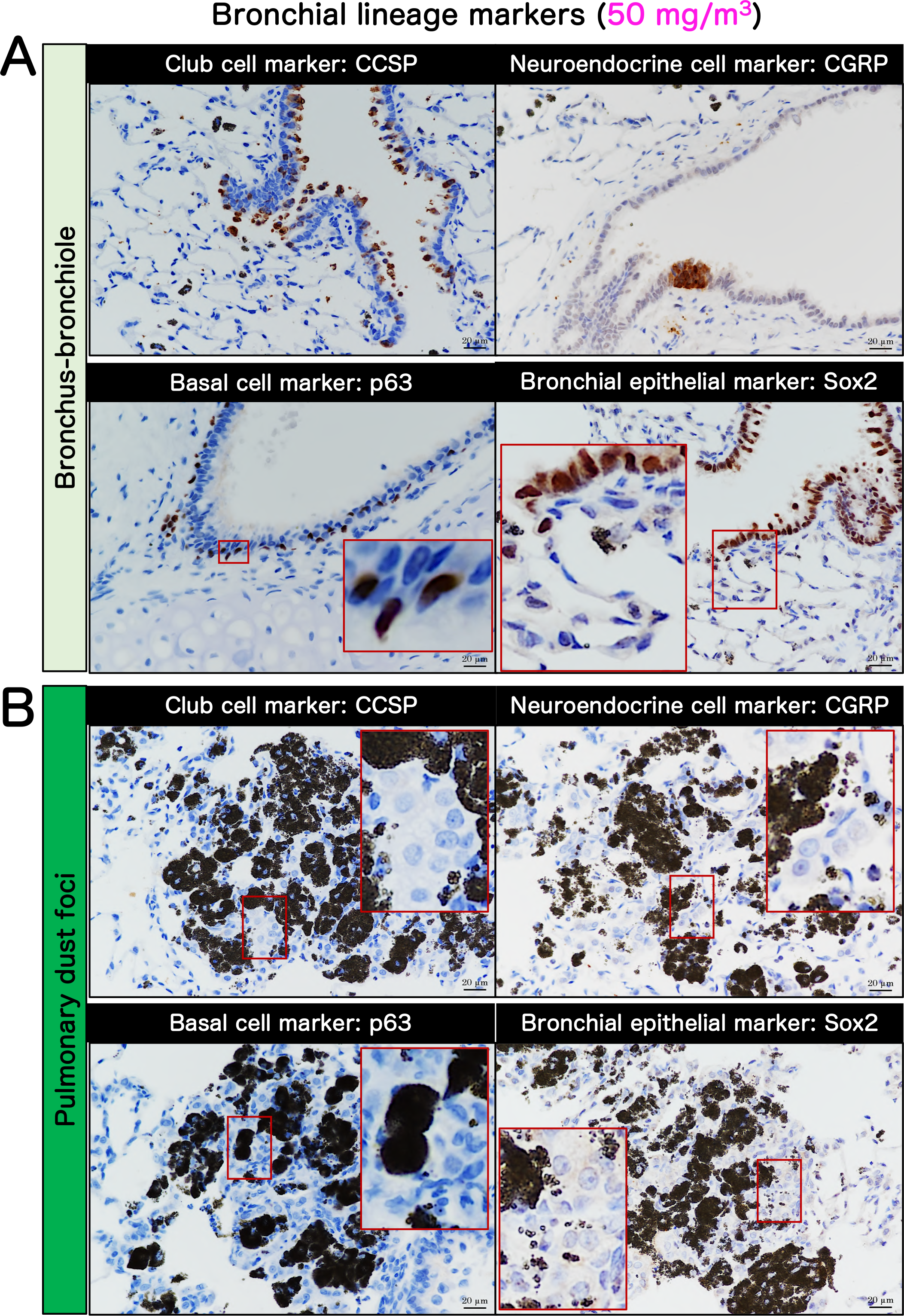
Immunohistochemistrical characteristics of bronchial lineage markers in tissue surrounding a lesion and pulmonary dust foci. Representative immunohistochemical staining images of the club cell marker club cell secretory protein (CCSP), neuroendocrine cell marker calcitonin gene-related peptide (CGRP), basal cell marker p63, bronchial epithelial lineage marker SRY-Box Transcription Factor 2 (Sox2) in the bronchus-bronchiole (A) and in pulmonary dust foci (B).

**Additional file 8: Figure S8.**
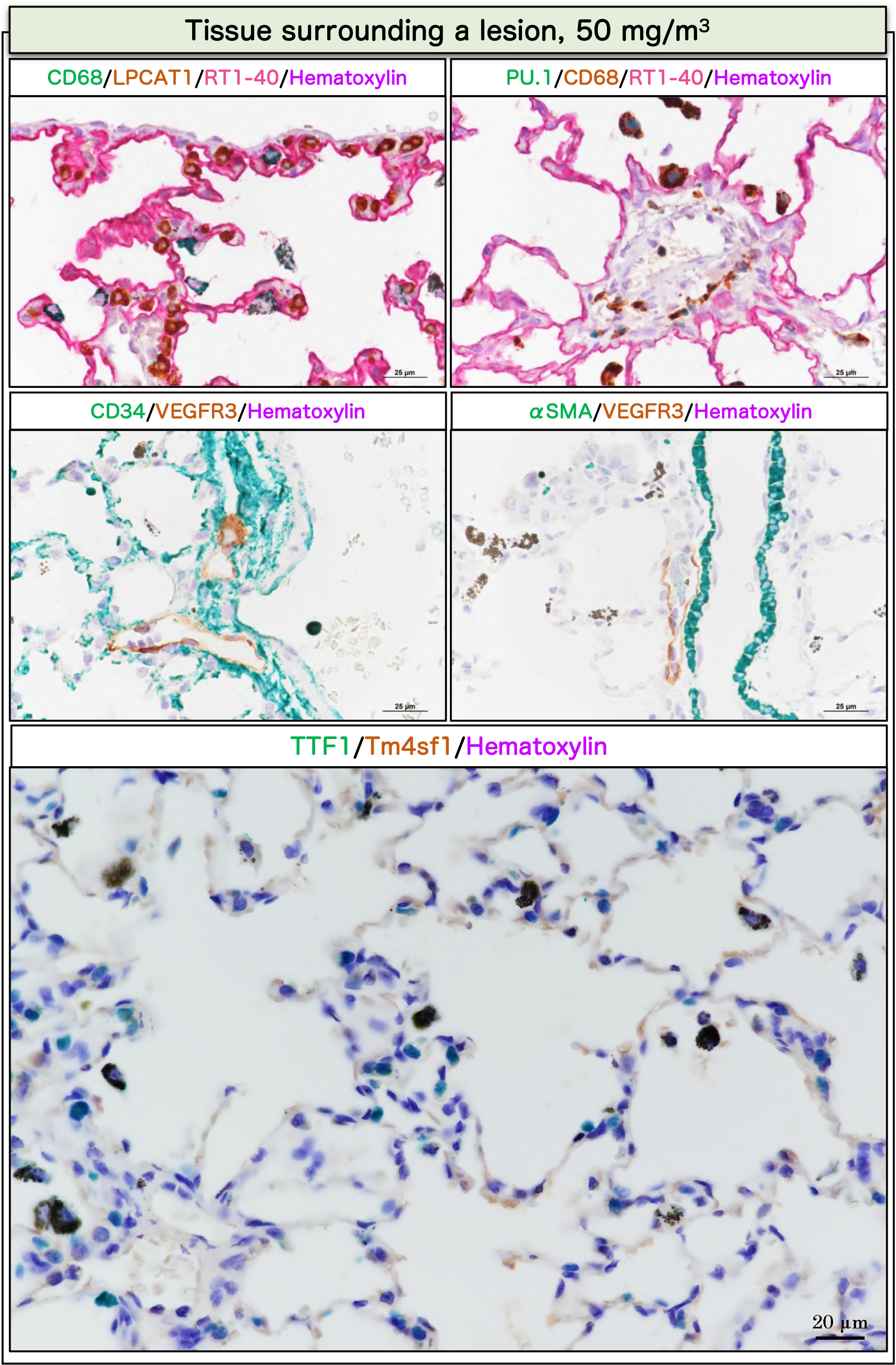
Immunohistochemical characteristics in tissue surrounding a lesion in rat lungs after inhalation exposure to TiO_2_ NP (50 mg/m^3^): same rat as in figure 9. Representative images of staining sets similar to figure 9 in tissue surrounding a lesion (normal tissue).

**Additional file 9: Figure S9.**
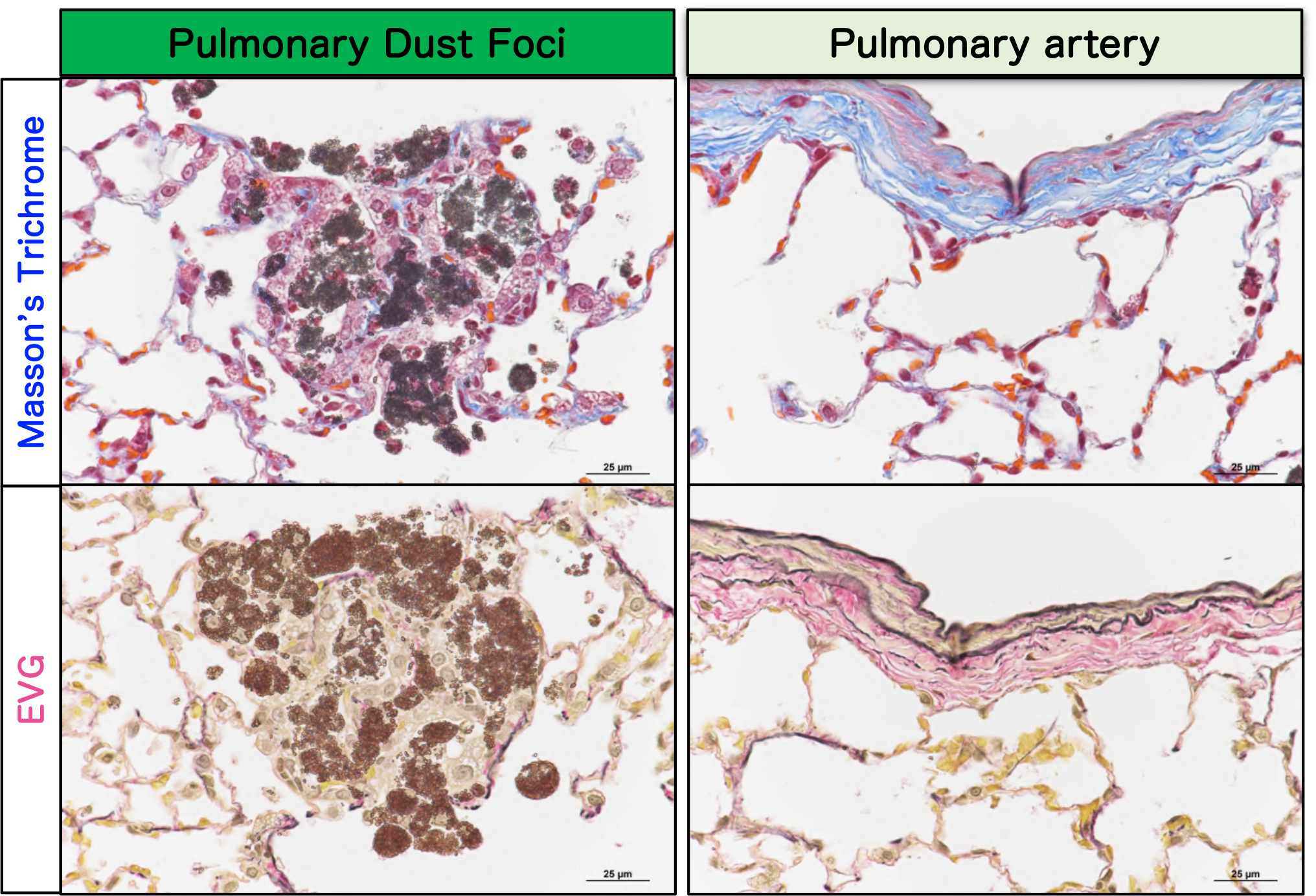
Representative microscopic photographs of Masson’s trichrome and EVG staining in pulmonary dust foci and pulmonary artery. Both stains were strongly positive in the arterial wall within the lung (right), but negative in the interstitium of the pulmonary dust foci (left). Abbreviations: EVG, Elastica Van Gieson.

**Additional file 10: Figure S10.**
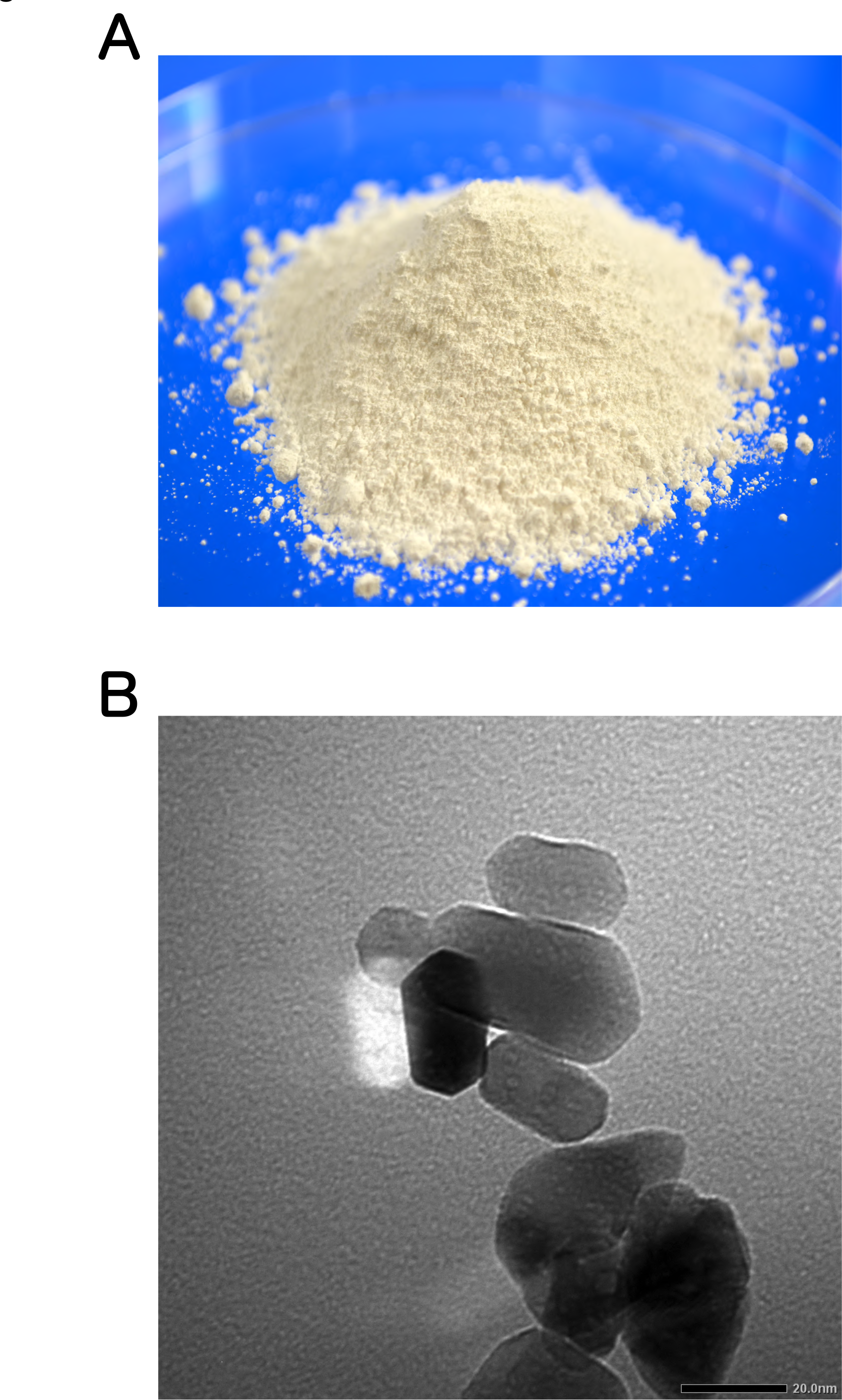
Representative macroscopic and SEM images of TiO_2_ NP. A: macroscopic image. B; SEM image.

**Additional file 11: Figure S11.**
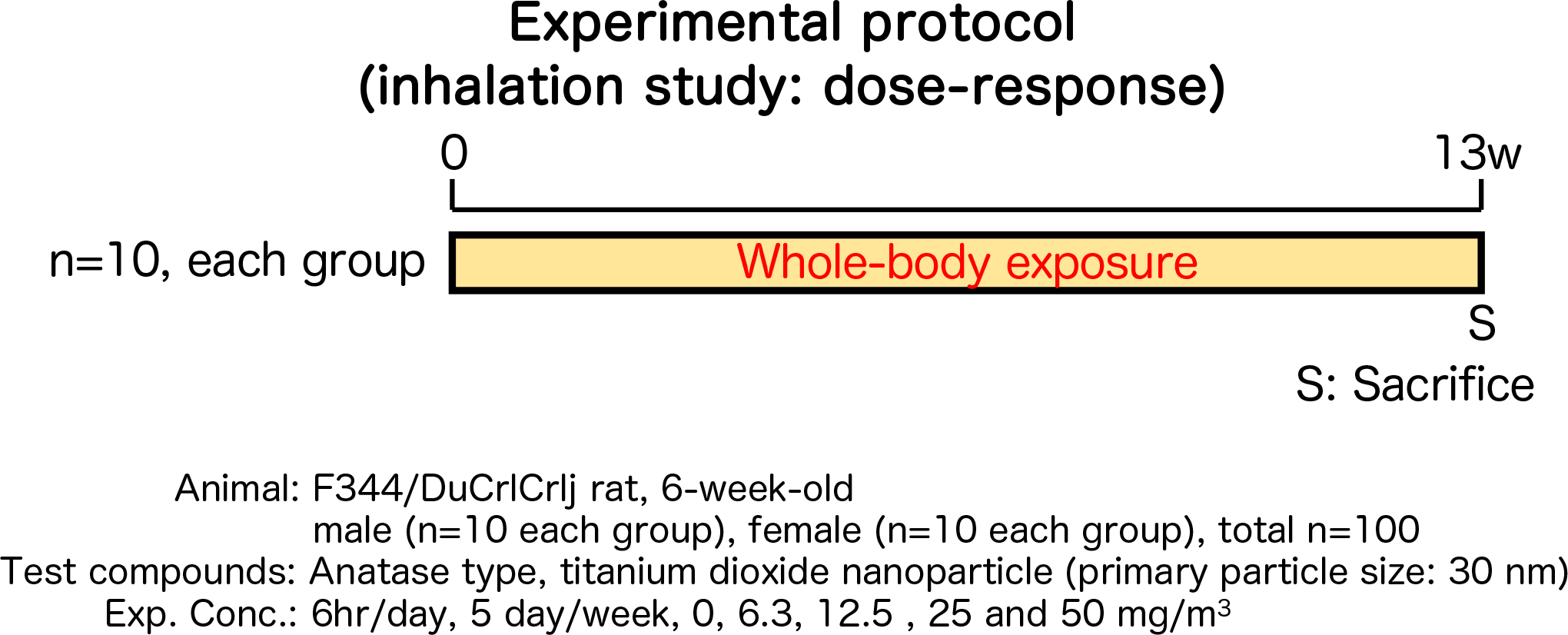
Design of animal experimental protocol in this study.

**Table S1.** Absolute organ weights observed in 13-week inhalation exposure study.

| Exposure concentration (mg/m<sup>3</sup>)<br>No. of Animals Examined | Male |  |  |  |  | Female |  |  |  |  |
| --- | --- | --- | --- | --- | --- | --- | --- | --- | --- | --- |
|  | 0 | 6.3 | 12.5 | 25 | 50 | 0 | 6.3 | 12.5 | 25 | 50 |
|  | 10 | 10 | 10 | 10 | 10 | 10 | 10 | 10 | 10 | 10 |
| Organ (g) |  |  |  |  |  |  |  |  |  |  |
| Thymus | 0.202±0.034 | 0.216±0.026 | 0.231±0.060 | 0.232±0.027 | 0.231±0.027 | 0.180±0.021 | 0.178±0.022 | 0.184±0.021 | 0.182±0.019 | 0.186±0.021 |
| Adrenals | 0.051±0.008 | 0.053±0.010 | 0.051±0.009 | 0.056±0.009 | 0.052±0.009 | 0.057±0.008 | 0.058±0.009 | 0.055±0.008 | 0.055±0.008 | 0.053±0.004 |
| Testes/Ovaries | 3.088±0.084 | 3.067±0.152 | 3.053±0.090 | 3.090±0.175 | 3.119±0.074 | 0.095±0.017 | 0.093±0.014 | 0.094±0.021 | 0.095±0.016 | 0.086±0.013 |
| Heart | 0.815±0.051 | 0.838±0.073 | 0.839±0.046 | 0.839±0.049 | 0.865±0.055 | 0.566±0.047 | 0.569±0.034 | 0.565±0.045 | 0.574±0.043 | 0.556±0.033 |
| Kidneys | 1.679±0.070 | 1.711±0.108 | 1.716±0.090 | 1.733±0.077 | 1.741±0.103 | 1.077±0.064 | 1.099±0.059 | 1.091±0.041 | 1.105±0.048 | 1.088±0.064 |
| Spleen | 0.556±0.043 | 0.552±0.035 | 0.578±0.028 | 0.584±0.030 | 0.578±0.047 | 0.352±0.029 | 0.360±0.025 | 0.376±0.038 | 0.368±0.028 | 0.362±0.025 |
| Liver | 6.563±0.439 | 6.628±0.542 | 6.786±0.383 | 6.823±0.453 | 6.873±0.474 | 3.756±0.253 | 3.723±0.243 | 3.818±0.263 | 3.823±0.309 | 3.737±0.212 |
| Brain | 1.871±0.050 | 1.870±0.027 | 1.904±0.047 | 1.901±0.023 | 1.888±0.036 | 1.762±0.061 | 1.746±0.024 | 1.757±0.070 | 1.751±0.060 | 1.760±0.039 |

**Table S2.** Relative organ weights observed in 13-week inhalation exposure study.

| Exposure concentration (mg/m<sup>3</sup>)<br>No. of Animals Examined | Male |  |  |  |  | Female |  |  |  |  |
| --- | --- | --- | --- | --- | --- | --- | --- | --- | --- | --- |
|  | 0 | 6.3 | 12.5 | 25 | 50 | 0 | 6.3 | 12.5 | 25 | 50 |
|  | 10 | 10 | 10 | 10 | 10 | 10 | 10 | 10 | 10 | 10 |
| Organ (%) |  |  |  |  |  |  |  |  |  |  |
| Thymus | 0.074±0.011 | 0.079±0.008 | 0.082±0.020 | 0.082±0.008 | 0.082±0.010 | 0.115±0.009 | 0.114±0.014 | 0.116±0.011 | 0.114±0.008 | 0.121±0.011 |
| Adrenals | 0.019±0.002 | 0.019±0.004 | 0.018±0.004 | 0.020±0.003 | 0.019±0.004 | 0.037±0.005 | 0.037±0.006 | 0.034±0.004 | 0.035±0.006 | 0.034±0.003 |
| Testes/Ovaries | 1.142±0.075 | 1.118±0.079 | 1.087±0.050 | 1.098±0.080 | 1.106±0.038 | 0.061±0.009 | 0.060±0.009 | 0.059±0.012 | 0.060±0.009 | 0.056±0.008 |
| Heart | 0.301±0.014 | 0.305±0.021 | 0.298±0.016 | 0.298±0.013 | 0.306±0.010 | 0.362±0.022 | 0.363±0.022 | 0.355±0.024 | 0.363±0.016 | 0.361±0.018 |
| Kidneys | 0.620±0.033 | 0.622±0.018 | 0.611±0.026 | 0.615±0.029 | 0.617±0.028 | 0.689±0.024 | 0.701±0.036 | 0.685±0.028 | 0.699±0.040 | 0.706±0.030 |
| Spleen | 0.205±0.011 | 0.201±0.004 | 0.206±0.014 | 0.207±0.009 | 0.205±0.010 | 0.225±0.013 | 0.229±0.011 | 0.235±0.016 | 0.232±0.009 | 0.235±0.012 |
| Liver | 2.419±0.069 | 2.407±0.047 | 2.413±0.095 | 2.417±0.076 | 2.433±0.080 | 2.403±0.121 | 2.373±0.054 | 2.394±0.098 | 2.412±0.102 | 2.426±0.100 |
| Brain | 0.693±0.056 | 0.682±0.042 | 0.678±0.039 | 0.675±0.029 | 0.670±0.028 | 1.128±0.040 | 1.116±0.063 | 1.103±0.047 | 1.108±0.057 | 1.144±0.046 |

**Table S3.** Blood-hematologic data observed in 13-week inhalation exposure study.

| Exposure concentration (mg/m<sup>3</sup>)<br>No. of Animals Examined | Male |  |  |  |  | Female |  |  |  |  |
| --- | --- | --- | --- | --- | --- | --- | --- | --- | --- | --- |
|  | 0<br>5 | 6.3<br>5 | 12.5<br>5 | 25<br>5 | 50<br>5 | 0<br>5 | 6.3<br>5 | 12.5<br>5 | 25<br>5 | 50<br>5 |
| Hematology |  |  |  |  |  |  |  |  |  |  |
| RBCs (10<sup>6</sup>/μl) | 9.35±0.20 | 9.18±0.28 | 9.30±0.17 | 9.37±0.21 | 9.34±0.09 | 8.61±0.33 | 8.63±0.17 | 8.54±0.17 | 8.31±0.21 | 8.69±0.47 |
| Hemoglobin (g/dl) | 15.4±0.2 | 15.2±0.6 | 15.3±0.4 | 15.4±0.3 | 15.3±0.3 | 15.2±0.6 | 15.2±0.2 | 15.0±0.4 | 14.8±0.3 | 15.5±1.0 |
| Hmatocrit (%) | 45.4±0.6 | 44.8±0.9 | 44.9±0.8 | 45.5±1.2 | 45.1±0.6 | 43.9±1.7 | 44.3±1.0 | 44.0±0.8 | 43.0±1.1 | 44.4±2.2 |
| MCV (fl) | 48.5±0.6 | 48.9±0.6 | 48.3±0.3 | 48.6±0.5 | 48.3±0.4 | 51.0±0.5 | 51.3±0.3 | 51.5±0.3 | 51.7±0.1* | 51.1±0.3 |
| MCH (pg) | 16.5±0.2 | 16.5±0.2 | 16.4±0.2 | 16.5±0.2 | 16.3±0.2 | 17.7±0.2 | 17.6±0.3 | 17.5±0.2 | 17.7±0.2 | 17.8±0.3 |
| MCHC (g/dl) | 33.9±0.5 | 33.9±0.7 | 34.0±0.4 | 33.9±0.6 | 33.9±0.7 | 34.6±0.7 | 34.3±0.7 | 34.0±0.5 | 34.3±0.5 | 34.8±0.6 |
| Platelet (10<sup>3</sup>/μl) | 762±31 | 719±29 | 751±42 | 773±46 | 756±65 | 793±48 | 766±42 | 799±48 | 788±43 | 750±62 |
| Reticulocyte (%) | 2.1±0.1 | 2.2±0.3 | 2.0±0.1 | 2.2±0.1 | 2.0±0.2 | 1.7±0.1 | 1.9±0.2 | 2.0±0.2 | 1.9±0.3 | 1.9±0.2 |
| Prothrombin Time (sec) | 12.3±0.3 | 12.3±0.4 | 12.4±0.1 | 12.2±0.1 | 12.3±0.2 | 11.9±0.2 | 12.3±0.3 | 12.2±0.3 | 12.0±0.5 | 12.1±0.5 |
| APTT (sec) | 16.6±0.6 | 17.0±0.8 | 17.6±0.6 | 17.3±0.5 | 17.8±0.6 | 13.2±0.9 | 13.6±1.1 | 14.2±0.6 | 13.3±0.7 | 12.6±1.3 |
| WBC (10<sup>3</sup>/μl) | 4.50±0.88 | 3.54±0.49 | 3.64±0.33 | 4.01±0.74 | 3.69±0.56 | 2.07±0.56 | 1.80±0.50 | 1.85±0.43 | 1.98±0.30 | 2.39±0.84 |
| Differential WBC (%) |  |  |  |  |  |  |  |  |  |  |
| Neutrophil | 31.4±5.2 | 34.6±5.0 | 30.3±4.2 | 31.7±4.6 | 31.9±0.7 | 25.0±6.2 | 29.5±7.0 | 26.8±3.3 | 29.3±5.0 | 30.1±4.3 |
| Lymphocyte | 64.6±5.0 | 61.6±5.0 | 65.4±4.0 | 64.2±4.9 | 63.6±0.8 | 70.4±6.6 | 65.7±6.6 | 68.6±3.1 | 65.8±5.3 | 65.4±4.2 |
| Monocyte | 1.7±0.2 | 1.6±0.4 | 1.7±0.2 | 1.5±0.3 | 1.6±0.5 | 2.0±0.6 | 1.9±0.3 | 1.9±0.4 | 2.1±0.4 | 1.7±0.3 |
| Eosinophil | 1.1±0.2 | 1.4±0.1 | 1.4±0.2* | 1.5±0.3* | 1.8±0.2** | 1.1±0.2 | 1.5±0.3 | 1.4±0.3 | 1.7±0.5 | 1.8±0.5 |
| Basophil | 0.1±0.1 | 0.1±0.1 | 0.1±0.1 | 0.1±0.1 | 0.1±0.1 | 0.3±0.1 | 0.3±0.2 | 0.1±0.1 | 0.1±0.1 | 0.2±0.2 |
| Other | 1.1±0.2 | 0.6±0.3 | 1.0±0.2 | 0.9±0.2 | 1.0±0.3 | 1.1±0.1 | 1.0±0.7 | 1.3±0.8 | 0.9±0.1 | 0.9±0.4 |

**Table S4.** Blood-biochemistry data observed in 13-week inhalation exposure study.

| Exposure concentration (mg/m<sup>3</sup>)<br>No. of Animals Examined | Male |  |  |  |  | Female |  |  |  |  |
| --- | --- | --- | --- | --- | --- | --- | --- | --- | --- | --- |
|  | 0 | 6.3 | 12.5 | 25 | 50 | 0 | 6.3 | 12.5 | 25 | 50 |
|  | 5 | 5 | 5 | 5 | 5 | 5 | 5 | 5 | 5 | 5 |
| Biochemistry |  |  |  |  |  |  |  |  |  |  |
| Total protein (g/dl) | 6.1±0.1 | 6.1±0.2 | 6.1±0.1 | 6.2±0.1 | 6.1±0.2 | 6.1±0.1 | 6.2±0.1 | 6.2±0.1 | 6.1±0.2 | 6.1±0.2 |
| Albumin (g/dl) | 3.3±0.1 | 3.3±0.2 | 3.4±0.1 | 3.3±0.0 | 3.3±0.1 | 3.4±0.1 | 3.5±0.1 | 3.5±0.1 | 3.4±0.1 | 3.4±0.1 |
| A/G ratio | 1.2±0.0 | 1.2±0.1 | 1.2±0.0 | 1.2±0.0 | 1.2±0.1 | 1.2±0.1 | 1.3±0.1 | 1.3±0.1 | 1.2±0.1 | 1.3±0.0 |
| T-Bilirubin (mg/dl) | 0.03±0.00 | 0.03±0.01 | 0.03±0.01 | 0.03±0.00 | 0.03±0.01 | 0.03±0.01 | 0.04±0.01 | 0.03±0.01 | 0.04±0.01 | 0.03±0.01 |
| Glucose (mg/dl) | 202±11 | 218±13 | 214±21 | 210±15 | 215±17 | 134± 6 | 137±10 | 124± 9 | 131± 7 | 143± 4 |
| T-Cholesterol (mg/dl) | 71± 6 | 68± 3 | 70± 4 | 70± 4 | 67± 5 | 79± 5 | 79± 9 | 83± 4 | 78± 9 | 84± 7 |
| Triglyceride (mg/dl) | 62±15 | 54± 7 | 53±13 | 49± 9 | 59± 5 | 18± 3 | 15± 6 | 12± 2 | 16± 6 | 23± 5 |
| Phospholipid (mg/dl) | 117± 7 | 116± 5 | 115± 6 | 115± 4 | 114± 5 | 145± 9 | 140±17 | 149± 6 | 139±15 | 149±16 |
| AST (U/l) | 102±27 | 161±45 | 126±53 | 111±37 | 90±18 | 60± 5 | 73± 9** | 63± 4 | 63± 5 | 71± 4* |
| ALT (U/l) | 59±12 | 78±20 | 66±19 | 57± 9 | 53±10 | 32± 8 | 40± 9 | 28± 4 | 31± 3 | 34± 5 |
| LDH (U/l) | 122±25 | 179±51 | 131±54 | 106±40 | 90±18 | 62±10 | 73±15 | 82±26 | 93±32 | 121±38** |
| ALP (U/l) | 360±23 | 373±22 | 400±45 | 390±31 | 373±22 | 305±31 | 307±15 | 293±28 | 303±44 | 326±24 |
| G-GTP (U/l) | 0.7±0.1 | 0.6±0.1 | 0.7±0.2 | 0.7±0.1 | 0.6±0.1 | 0.9±0.1 | 1.1±0.2 | 1.0±0.2 | 1.0±0.2 | 1.2±0.3 |
| CK (U/l) | 105±14 | 119±38 | 98±16 | 105±15 | 101±13 | 88±13 | 94± 7 | 101±14 | 114±25 | 138±44 |
| Ureanitrogen (mg/dl) | 20.6±1.1 | 20.1±1.3 | 21.5±1.2 | 22.1±1.2 | 20.1±1.7 | 19.0±0.8 | 21.9±1.8* | 20.1±1.6 | 21.0±1.5 | 23.7±0.8** |
| Creatinine | 0.32±0.03 | 0.33±0.03 | 0.36±0.02 | 0.35±0.04 | 0.33±0.03 | 0.34±0.02 | 0.34±0.03 | 0.35±0.03 | 0.36±0.02 | 0.34±0.04 |
| Sodium (mEq/l) | 142± 1 | 142± 1 | 142± 1 | 142± 0 | 142± 0 | 143± 1 | 143± 1 | 144± 1 | 144± 1 | 143± 1 |
| Potassium (mEq/l) | 3.2±0.1 | 3.1±0.1 | 3.0±0.1 | 3.0±0.1 | 3.0±0.2 | 3.1±0.1 | 3.0±0.1 | 3.1±0.2 | 3.1±0.2 | 3.2±0.2 |
| Chloride (mEq/l) | 103± 1 | 103± 1 | 102± 0 | 102± 1 | 102± 1 | 104± 1 | 105± 2 | 105± 1 | 105± 1 | 104± 1 |
| Calcium (mg/dl) | 10.1±0.1 | 9.9±0.1 | 10.0±0.2 | 9.9±0.1 | 10.0±0.2 | 9.9±0.1 | 9.8±0.3 | 9.8±0.1 | 10.0±0.2 | 9.9±0.3 |
| Inorganic phosphrus (mg/dl) | 5.6±0.3 | 5.0±0.4 | 5.2±0.3 | 5.3±0.2 | 5.1±0.4 | 5.5±0.6 | 4.6±0.7 | 5.0±0.4 | 6.0±0.9 | 6.2±1.3 |

**Table S5.**
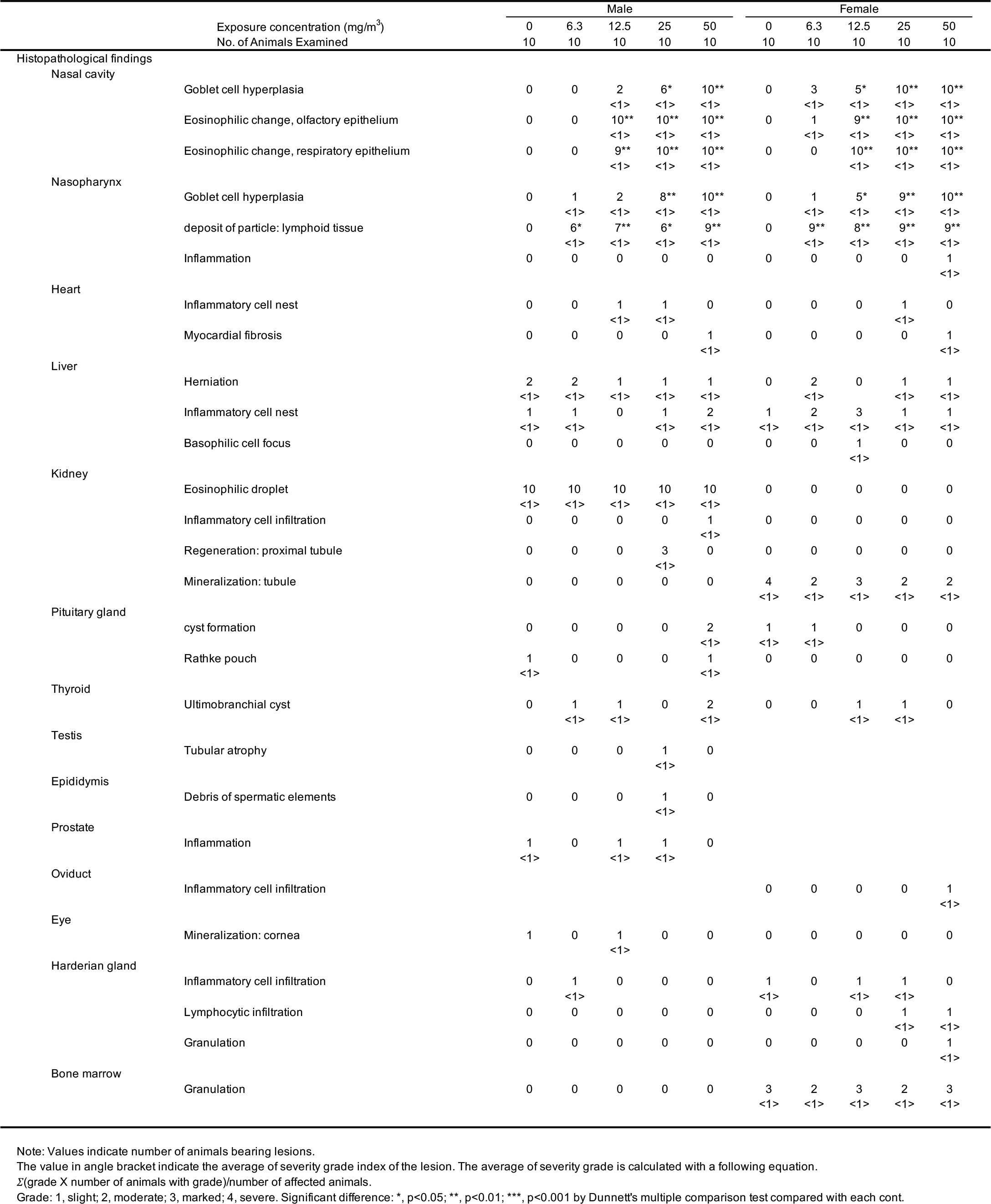
Histopathological findings excluding lung and mediastinal lymph node observed in 13-week inhalation exposure study.

**Table S6.** The summary of the effect of inhalation exposure to TiO_2_ (in this study) and ultrafine TiO_2_ particle (P25) on LDH activity.

| Test compounds | 13-week study | chamber condition |  | LDH in BALF |  |  |
| --- | --- | --- | --- | --- | --- | --- |
|  |  | MMAD<br>(σg)<br>(μm) | Exposure<br>concentration<br>(mg/m<sup>3</sup>) | average ± SD<br>(U/l) |  | Fold<br>change |
| TiO<sub>2</sub> NP (in-house, in this study) |  |  |  |  |  |  |
|  | male F344/DuCrI CrIj rat |  | 0 | 24 ± | 4.7 | - |
|  |  | 0.9 (2.1) | 6.3 | 28 ± | 3.8 | 1.2 |
|  |  | 0.9 (2.1) | 12.5 | 30 ± | 2.3 | 1.3 |
|  |  | 0.9 (2.1) | 25 | 33 ± | 1.4 | 1.4 |
|  |  | 1.0 (2.1) | 50 | 60 ± | 2.7 | 2.5 |
|  | female F344/DuCrI CrIj rat |  | 0 | 30 ± | 2.8 | - |
|  |  | 0.9 (2.1) | 6.3 | 36 ± | 2.4 | 1.2 |
|  |  | 0.9 (2.1) | 12.5 | 38 ± | 4.7 | 1.2 |
|  |  | 0.9 (2.1) | 25 | 42 ± | 2.1 | 1.4 |
|  |  | 1.0 (2.1) | 50 | 108 ± | 16.2 | 3.6 |
| TiO<sub>2</sub>, ultrafine (Bermudez *et al.*) |  |  |  |  |  |  |
|  | female CDF(F344)/CrI BR rat |  | 0 | 24 ± | 2.0 | - |
|  |  | 1.4 (2.6) | 0.5 | 26 ± | 2.0 | 1.1 |
|  |  | 1.4 (2.6) | 2 | 29 ± | 7.0 | 1.2 |
|  |  | 1.4 (2.6) | 10 | 122 ± | 18.0 | 5.1 |

**Table S7.**
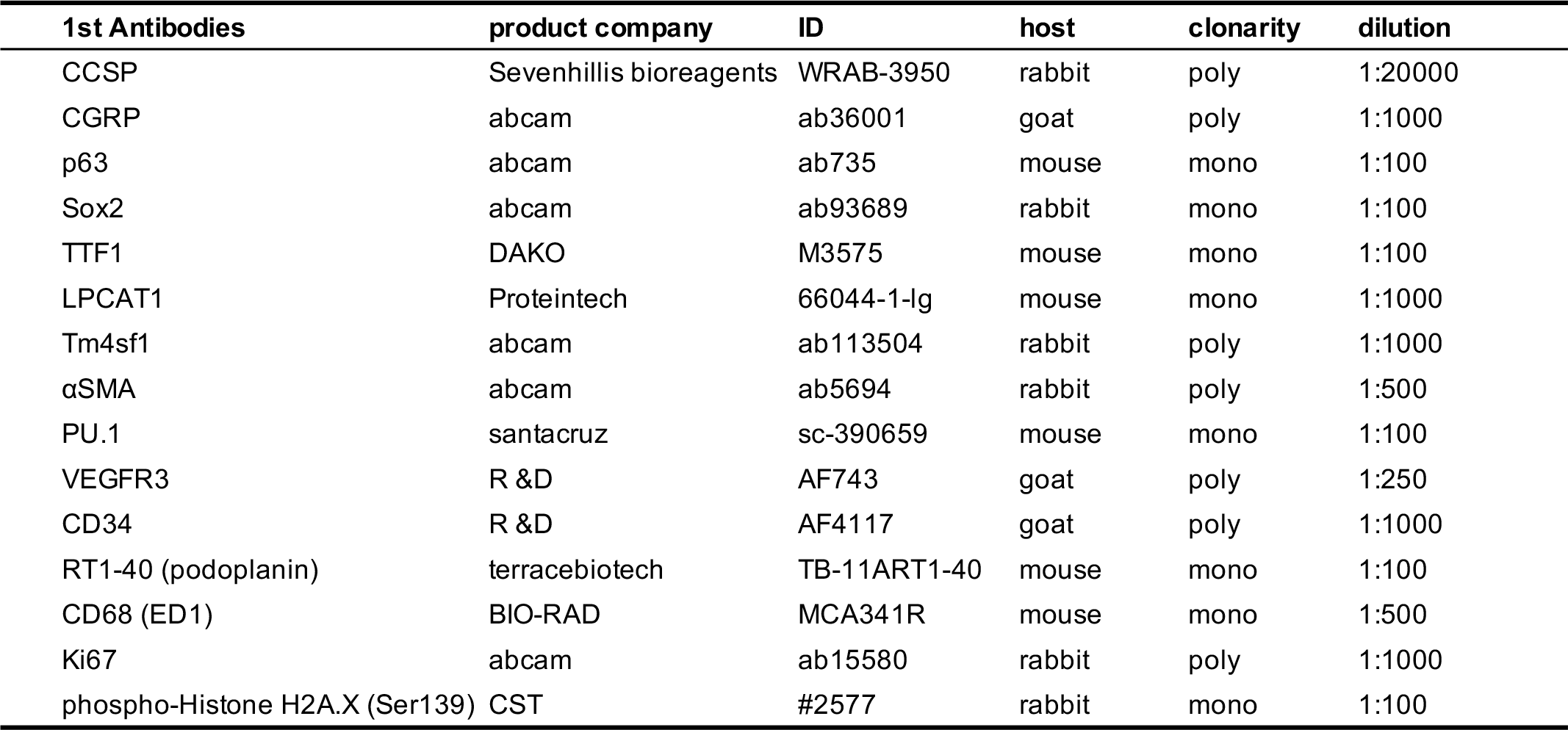
List of primary antibodies used in this study.

| 1st Antibodies | product company | ID | host | clonarity | dilution |
| --- | --- | --- | --- | --- | --- |
| CCSP | Sevenhillis bioreagents | WRAB-3950 | rabbit | poly | 1:20000 |
| CGRP | abcam | ab36001 | goat | poly | 1:1000 |
| p63 | abcam | ab735 | mouse | mono | 1:100 |
| Sox2 | abcam | ab93689 | rabbit | mono | 1:100 |
| TTF1 | DAKO | M3575 | mouse | mono | 1:100 |
| LPCAT1 | Proteintech | 66044-1-Ig | mouse | mono | 1:1000 |
| Tm4sf1 | abcam | ab113504 | rabbit | poly | 1:1000 |
| αSMA | abcam | ab5694 | rabbit | poly | 1:500 |
| PU.1 | santacruz | sc-390659 | mouse | mono | 1:100 |
| VEGFR3 | R &D | AF743 | goat | poly | 1:250 |
| CD34 | R &D | AF4117 | goat | poly | 1:1000 |
| RT1-40 (podoplanin) | terracebiotech | TB-11ART1-40 | mouse | mono | 1:100 |
| CD68 (ED1) | BIO-RAD | MCA341R | mouse | mono | 1:500 |
| Ki67 | abcam | ab15580 | rabbit | poly | 1:1000 |
| phospho-Histone H2A.X (Ser139) | CST | #2577 | rabbit | mono | 1:100 |

## Notes

### Competing Interest Statement

The authors have declared no competing interest.

